# Evaluating a structured expert elicitation approach for adaptive conservation: Lessons from five years in practice

**DOI:** 10.1101/2025.11.08.687399

**Authors:** Helen J Mayfield, James Brazill-Boast, Mick Andren, Michiala Bowen, Adam Fawcett, Trent Forge, Luke Foster, Ross Goldingay, Paul Hillier, Meagan Hinds, Simon Lee, Erica Mahon, Martine Maron, Doug Mills, Thomas Rowell, Stephanie Stuart, Caren Taylor, Grant Webster, Nicole Hansen

## Abstract

Threatened species management relies on *Ex ante* estimates of species’ responses to different interventions to generate meaningful predictions. Structured expert elicitation is often used to generate these estimates, but comparisons of these expert-predicted outcomes with observed results are rare. This study aims to evaluate the utility of expert elicitation for adaptive management in the New South Wales Saving our Species (SoS) program in Australia by revisiting six species management plans that were generated from bespoke structured elicitation guidelines five years prior. Each species’ management plan included a defined scope, conceptual model, monitoring indicators and estimated response to management curves under different scenarios. Experts reviewed the conceptual models after five years of management and monitoring and compared the predicted response to management with observed monitoring data. In three of the six case studies, observed outcomes closely matched predictions. Where predictions diverged, factors such as unanticipated new threats and unexpected responses to interventions contributed to discrepancies. However, in all cases, the structured approach provided a clear logic for planning, enabling managers to systematically refine their understanding. The conceptual models and response curves proved valuable for collaboration, communication, and generating hypotheses for unexpected results. This work demonstrates the value of the bespoke guidelines in supporting adaptive management processes, strengthening the knowledge base for threatened species conservation while improving alignment between predictions and real-world outcomes.

## Introduction

An increasing number of species around the world require urgent, ongoing management to prevent their extinction (Díaz et al. 2019, Gumbs et al. 2024, Lanzas et al. 2024). Those working to prevent the loss of the world’s biodiversity do so with limited resources and an imperfect understanding of threatening processes and the factors that determine how species or ecosystems respond to management interventions. To prevent extinction, managers are often required to rapidly implement practical interventions and monitoring programs despite this uncertainty, while still considering complex environmental factors and social, cultural, and political interactions. In these instances, adopting an adaptive management approach that is flexible yet scientifically robust is critical as it encourages conservation practitioners to learn progressively and change strategies to exploit evolving knowledge or other opportunities (Walters et al. 1990, Kingsford et al. 2011).

Adaptive management is a systematic approach to improving resource management by learning from, and responding to, the outcomes of successive interventions (Gregory 2012). It combines project design, management and monitoring with a framework for testing assumptions; practitioners adapt their management plans if the system does not respond to the interventions as expected (Margoluis 1998). As part of a structured decision-making process (Gregory 2012), adaptive management can help managers specifically address significant uncertainty surrounding the most effective or efficient course of action (Hemming et al. 2022). It requires a shared understanding of the problem to form the basis of hypotheses about how a species will respond to alternative management interventions, as well as a set of relevant, consistent ecological indicators that are both feasible to monitor and sensitive enough to provide reliable data to test those hypotheses (Lindenmayer et al. 2020). Finally, in contrast to passive surveillance, a key requirement for adaptive management is having some *a priori* estimate of what is expected, to allow for comparison with a competing hypothesis or counterfactual (Nichols et al. 2006).

Limited resources, and the urgency of action given often widespread and severe threats, commonly results in a lack of carefully considered hypotheses, appropriate control sites or counterfactual models. Without these, it becomes impossible to understand the effectiveness of management interventions. In the absence of a control site, or sufficient data and resources to model counterfactual scenarios, expert knowledge elicited in a structured fashion is often a feasible alternative (Martin et al. 2012, Burgman 2016, Hemming et al. 2018). As well as providing a counterfactual for guiding adaptive management, expert-elicited approaches based on conceptual modelling (Schwartz et al. 2012, Mayfield et al. 2020) are valuable for helping practitioners better understand the complexities of their conservation projects, such as the multitude of factors operating at a site that are likely to impact species. By supporting practitioners to document the pathways between their management actions and forecasted results, structured expert-elicited approaches can facilitate the design of robust interventions and selection appropriate indicators to measure project success.

The Saving our Species program (Brazill-Boast et al. 2018) is an initiative of the New South Wales (NSW) State Government in Australia that encapsulates a robust yet practical approach to science-based management of threatened species and ecosystems. The program aims to secure as many species as possible in the wild in NSW for the next 100 years. Where possible, monitoring data collected from these projects are used to inform complex models to guide further management decisions. However, in many cases these data or resources are not available, particularly for cryptic or newly described species. In response, the Saving our Species program developed practical guidelines to assist managers to formalise their own expert knowledge into conceptual models and estimates of how a species is expected to respond to management over time, as well as making explicit the assumptions about the mechanisms driving that response.

Expert elicitation, such as that used in the NSW Saving our Species program, is becoming a common tool in environmental management (Adams-Hosking et al. 2016, Kleitou et al. 2021, Camus et al. 2022, Mayfield et al. 2023). Despite often being used to generate long-range projections, sometimes for many years into the future, there is rarely an opportunity in conservation to compare the expert-predicted values with the observed outcomes of the management actions. Following five years of implementation and monitoring, the Saving our Species program provides a unique opportunity to compare expert-generated conceptual models and estimated response-to-management curves with the observed outcomes of conservation projects.

By revisiting existing management plans devised five years prior, as well as describing two recent examples, this study aims to evaluate the utility of the Saving our Species approach to support adaptive management. Specifically, we evaluate 1) how closely the estimated response to management aligned with the observed outcomes; 2) whether the inclusion of conceptual models helped interpret any deviations from the expected values; and 3) whether the process was considered practical by practitioners tasked with designing management plans for threatened species and ecosystems. Additionally, we describe the key lessons learnt during the review process that contribute to the growing literature on expert elicitation for conservation management.

## Methods

### Saving our Species guidelines for estimating a species’ response to management

The Saving our Species guidelines (Mayfield et al. 2019) were devised in 2018 to assist those tasked with devising management plans to formalise their knowledge, and the knowledge of other experts, into a quantifiable adaptive monitoring and management plan. The guidelines were developed via a collaborative process involving academics, Saving our Species project coordinators (assigned to design and coordinate the Saving our Species management of the species or ecosystem across all sites), and other species’ experts (Mayfield, Rhodes et al. 2019, Mayfield, Brazill-Boast et al. 2020). The process in the guidelines sets out five steps (Fig 1). The outputs at each step form key components of the five-year adaptive management plan required for all managed species and ecosystems under the program. *Step 1*, defining the scope, outlines the sites, threats and actions that are being considered in the estimates. A conceptual model (*Step 2*) is then generated that defines the assumptions used when designing management interventions. This model guides the selection of monitoring indicators in *Step 3*, covering both the threatened species or ecoregion as well as the threats being actively managed. For each indicator, experts are then tasked with estimating the expected management response for each indicator (*Step 4*) for both the planned and a counterfactual management scenario, and at different time steps. Finally, *Step 5* requires selecting a range of benchmark (target) values for each indicator that can be compared against to demonstrate if management actions have been successful. Although represented in Fig. 1 as a linear process, it is often necessary to revise the output from previous steps as new information becomes available.

**Fig. 1.**
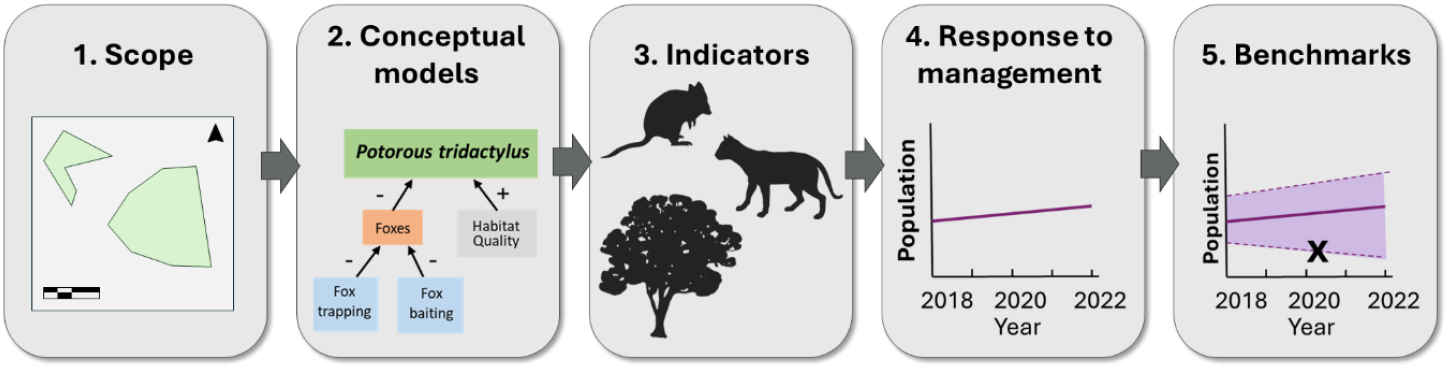
The five steps for the Saving our Species guidelines to estimating a species’ response to management

The guidelines also contain twelve real-world case studies of Saving our Species managed species, each including a conceptual model linking the proposed management actions to the species threats, and selected indicators for monitoring. Ten of these case studies also provide estimated response to management curves (response curves) for each indicator over time for both an expected and counterfactual scenario. Estimated response curves include the experts’ judgements on the lowest, highest and most likely values at five time points; 2016 (start of the Saving our Species program), 2017,2021, 2036 and 2116

### Case study review process

Six of the original twelve case studies from the Saving our Species guidelines were selected for review. Case studies were selected was based on availability of the current project coordinator, completeness of the original 2018 case study, and to represent a range of taxa. Conservation outcomes were not considered until after the case studies had been selected. The current project coordinator for each selected case study was invited to participate in the review and were encouraged to recommend any additional experts who had worked closely with the species over the previous five years.

Once the project coordinator had agreed to participate, the review team searched the internal Saving our Species management database for each case study to extract the planned and implemented management actions and monitoring data from 2017 to 2023. This included both qualitative and quantitative data for indicators relating to the species and threats being managed. Project coordinators were also able to provide additional data not yet entered in the system. For Objective 1, to compare the estimated and observed outcomes, we compared the results from the database, combined with any extra information provided by the project coordinators, with the original estimated response curves from the 2018 case studies. In instances where the indicators that were actually measured differed from those estimated in 2018, trends were still examined where the same attribute was measured in some other way, e.g. number of snakes vs occupancy for measuring the snake population.

For Objectives 2 and 3, relating to the interpretation of the results and usefulness of the process, semi-structured interviews were conducted in November and December of 2023 with project coordinators and the nominated experts, in person where possible. Prior to the interview, project coordinators were given the opportunity to review both the complete case study from 2018 and the monitoring summaries extracted from the management database. During the interviews, which lasted for between half an hour and 1.5 hours, project coordinators and their invited experts were asked to provide an overview of how closely the management interventions implemented over the last five years matched the planned interventions, and whether the species had responded as expected. They also worked through the conceptual model and discussed if, in their view, the assumptions used in 2018 were still supported. At the end of the interview, project coordinators and experts were asked if they had found the review process useful for informing the next phase of their project. After the initial interview, participants were invited to further reflect on discrepancies between the original estimates and the observed monitoring data and refine any updates to the conceptual models. Human ethics approval for conducting interviews was obtained from The University of Queensland under protocol 2023/HE001287.

### Evaluating practicality

To explore the practicality of the Saving our Species approach, two additional multi-species case studies were considered as part of the review. The first focused on the Mount Kaputar Snail and Slug Threatened Ecological Community, and the second was a combined case study for a specialist invertebrate species, the black grass-dart butterfly (*Ocybadistes knightorum*) and its food plant - Floyd’s grass (*Alexfloydia repens*). Both examples were developed in a series of separate workshops in 2022, where project coordinators worked with facilitators (HM, NH) to develop conceptual models, write narratives and select indicators. After having worked through the full process, the project coordinators were asked to reflect how useful they found it for developing and communicating their management plan, and whether any amendments were needed to accommodate the added complexity of multi-versus single-species planning. The accuracy of predictions was not examined for these case studies due to the lack of adequate implementation time.

## Results

### Management implementation

Project coordinators for all six species agreed to participate in the review process (Table 1), with all except for the long-nosed potoroo involving at least one expert who contributed to the original 2018 case study. Two major unforeseen events affected the Saving our Species program during the five-year period and are relevant to the interpretation of our results. The Black Summer bushfires in 2019-2020 burnt through over 5.4 million hectares of bushland (DPIE 2020) and had severe impact on many threatened species and ecoregions in the state (Ward et al. 2020). As well as the direct impacts on threatened species and their habitat (van Eeden 2020), this catastrophic event further stretched limited resources for planned management interventions. In addition, the COVID-19 pandemic caused disruptions in implementing both management and monitoring activities throughout the state. Widespread lockdowns and closure of state borders within Australia in 2020and 2021 also created unique and unforeseen circumstances for those species where human disturbance is considered a major threat (Stenhouse et al. 2022, Tucker et al. 2023).

**Table 1.** Case studies included in the 2024 Saving our Species guidelines (SoS) review. Mt Kaputar threated ecological community and black grass-dart butterfly/Floyd’s grass are additional case studies not included in the original Saving our Species guidelines.

| Case study | Taxon | Number of experts in review | Number of sites included in review | Management actions carried out |
| --- | --- | --- | --- | --- |
| Long-nosed potoroo (*Potorous tridactylus*) | Mammal | 1 | 1 | Partial |
| Large bent-winged bat (*Miniopterus schreibersii oceanensis*) | Mammal | 1 | 1 | Full |
| Mahoney’s toadlet (*Uperoleia mahonyi*) | Amphibian | 2 | N/A | N/A |
| Tall rusthood orchid (*Pterostylis chaetophora*) | Plant | 1 | 1 | Partial |
| Caley’s grevillea (*Grevillea caleyi*) | Plant | 1 | 1 | Partial |
| Broad-headed snake (*Hoplocephalus bungaroides*) | Reptile | 5 | 4 | Partial |
| Mt Kaputar threatened ecological community | Ecological Community | 1 | N/A | N/A |
| Black grass-dart butterfly (*Ocybadistes knightorum*) & Floyd’s grass (*Alexfloydia repens*) | Invertebrate and Plant | 1 | N/A | N/A |

Implementation of management plans between 2017 and 2023, was varied. Only the large bent-winged bat project was able to fully implement their planned management actions, noting that emerging threats such as white-nose fungus and nearby windfarm developments remained of concern for this species. Three projects — Caley’s grevillea, the broad-headed snake, and the long-nosed potoroo — were able to implement most of the planned management action but were hindered by factors such as external land management decisions and resource constraints.

Management of the tall rusthood orchid faced significant challenges, including restricted site access during COVID-19, program defunding, new land developments overlapping its habitat, and herbivory by birds consuming underground tubers. Mahoney’s toadlet, a data-deficient species, had no predefined management actions and was instead the focus of intensive research efforts. Full case study review details are provided in Supplementary material S1.

### Monitoring data

Two species (long-nosed potoroo and large bent-winged bat) had at least partial monitoring data for the same indicators selected for that species in 2018 (Table *2*). For the remaining four, the indicators measured differed from those nominated in 2018. Monitoring data for the tall rusthood orchid changed from abundance-based to a grid-based occupancy measure in 2021, as this was considered more appropriate for a cryptic species that may not be visible above ground each year. The broad-headed snake was measured using site occupancy, rather than abundance. Participants reflected that this represented a difference between sites in the suitability of using capture-recapture techniques. For Caley’s grevillea, as fire did not occur at the site during the monitoring period, suitable habitat area (in hectares) was used as a proxy instead of post-fire seedling abundance to compare the response to management. For Mahoney’s toadlet, the planned monitoring was refined from number of adults and tadpoles per site to the number of calling males, and presence/absence per site measured via eDNA.

As per the guidelines (Mayfield, Rhodes et al. 2019), the threats being managed under each plan were also monitored (Table 2). Additional threats added to the original monitoring plan included occupancy of dingoes for the long-nosed potoroo, frequency of human disturbance at caves for the large bent-winged bat, and estimated proportion of snakes lost to poaching for the broad-headed snake. For Caley’s grevillea, whether or not each hectare received fire management was added as a qualitative threat indicator. Disturbance monitoring was expanding to include including habitat disturbance from any recreational use, rather than focusing only on damage from illegally created tracks. In some case studies, indicators were removed or not included in the final monitoring.

**Table 2.** Attributes for six example case studies from the Saving our Species guidelines along with their suggested indicators from 2018 and the actual measured indicators. Not all indicators listed were measured for each site or year. Occupancy here is defined as the percent of camera traps at the site where the species is detected. Monitoring for Mahoney’s toadlet has not commenced.

| Species | Attribute | Suggested indicator | Measured/planned indicator |
| --- | --- | --- | --- |
| **Long-nosed potoroo** | Potoroo | Occupancy | Potoroos Occupancy |
|  | Foxes | Occupancy | Fox occupancy |
|  | Cats | Occupancy | Cat occupancy |
|  | Fire regime | Habitat under suitable fire regime (ha) | Fire activity (qualitative description of fire activity) |
|  | Dingoes | - | Dingo occupancy |
| **Large bent-winged bat** | Bats | Number of bats in cave | Number of bats in cave |
|  | Roost access | Proportion of roost entrance covered by weeds | Proportion of roost entrance covered by weeds |
|  | Cats | Nights per year with cats observed | Number of cats detected during surveys |
|  | Disturbance | - | Disturbance at caves from people (y/n) |
| **Mahoney's toadlet** | Frogs | Number of adult frogs in population | Number of calling males<br>Presence/absence (eDNA) |
|  | Tadpoles | Number of tadpoles | - |
|  | Water quality | Water quality score | Water quality (pH and sediment levels) |
|  | Habitat damage | Habitat degradation by pigs (score) | - |
| **Tall rusthood orchid** | Orchids | Number of individuals | Number of individuals/<br>Grid-based occupancy model |
|  | Grazing damage | Proportion of flowerheads chewed | - |
|  | Weed | Weed cover (%) | Ground cover weeds (%)<br>Shrubby weeds (%) |
|  | Disturbance | - | - |
| **Caley's grevillea** | Grevillea | Post-fire seedling abundance | - |
|  | Habitat | Area (Ha) | Area (Ha) |
|  | Weed density | - | - |
|  | disturbance | Relative disturbance from illegal tracks | Disturbance |
|  | Fire | - | Fire management (qualitative) |
| **Broad-headed snake** | Snakes | Population size | Site occupancy (yes/no) |
|  | Habitat disturbance | Rocks disturbed (%) | Extent of bush rock habitat disturbed |
|  | Habitat | Rocks remaining (%) | - |
|  | Adult geckoes | Number of geckoes | - |
|  | Poaching | - | Likelihood of illegal collection |

Examples include removing habitat damage from pigs for Mahoney’s toadlet (no longer considered a threat) and weed density for Caley’s grevillea (lost access to site).

### Species outcomes

Quantitative data were available for each project, with the exception of Mahoney’s toadlet, which was still in a research phase of the project. Data for the broad-headed snake outcomes and threats were available for all four sites that were included within the scope of the original case study. Of the three sites included in the potoroo case study, data were only available for the Barrington Tops site. For the remaining three species, data were included that matched the site included in the case study.

The long-nosed potoroo occupancy (% of camera traps detecting potoroos over the season) at the Barrington Tops site (Fig. 2a) did not increase as much as expected, despite the expected observed decrease in the fox population due to successful fox management onsite in 2020. While the reported observed occupancy for potoroos was higher than the estimated values, this represents a difference in the scope, with the observed values representing the Barrington Tops site rather than the entire managed range of the species.

**Fig. 2.**
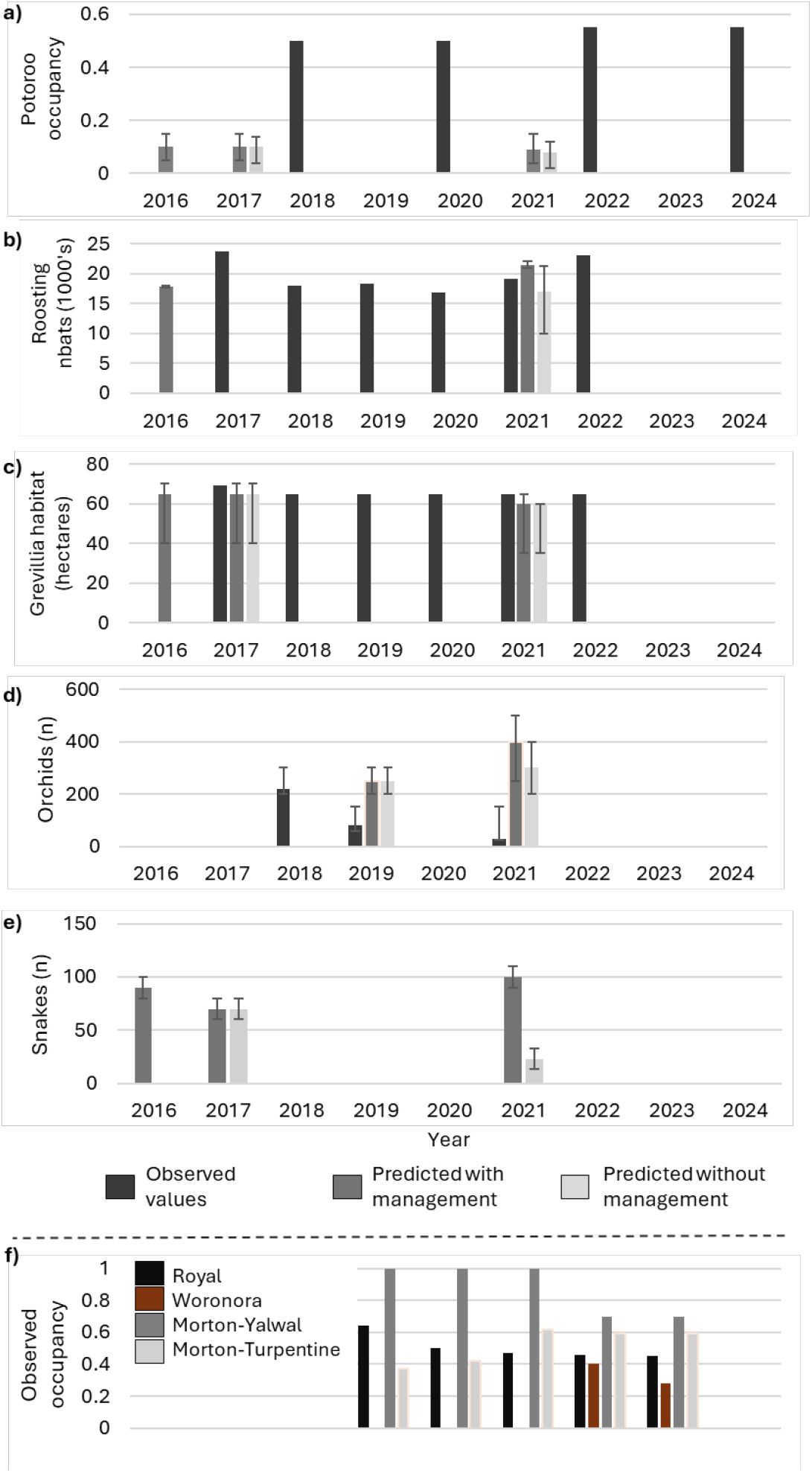
Observed and predicted values for a) long-nosed potoroo (Barrington Tops site), b) large bent-winged bat (Church cave site), c) tall rusthood orchid (Black Creek site), d) Caley’s grevillea, and e) broad-headed snake. Values (including confidence values) are given in supplementary Table S3.

The large bent-winged bat population at the Church cave site (Fig. 2b) responded as expected to the planned management actions (primarily clearing weeds from cave entrance), with all indicators remaining within the predicted limits. Likewise, the amount of suitable habitat recorded for Caley’s grevillea closely matched the predicted response curves despite recreational impacts still occurring in habitat areas (Fig. 2d). Tall rusthood data from the consistently monitored locations at the Black Creek site up until 2021 (Fig. 2c) show a substantial and alarming decrease in the number of orchids, below even that for the unmanaged scenario. After 2021, the discovery of orchids at additional onsite locations created an artificial upward trend (not shown). The broad-headed snake population responded well at Morton-turpentine in line with predictions and, with the exception of decreases caused by poaching events, remained stable at the other sites (Fig. 2e).

### Value of conceptual diagrams

Revisiting the conceptual diagrams for each of the 2018 case studies provided insights as to why the monitoring data did or did not accord with expectations and helped to identify practical ways to influence threats and conservation outcomes. For example, the broad-headed snake responded well, when considered in light of the previously unanticipated poaching events that depleted the population i.e. impacts of this poaching likely negating the expected occupancy gains from the implemented habitat improvements. The model was used to explore management interventions for reducing poaching, the prominent threat over the last five years, and how these actions might also reduce bush rock removal. Re-evaluating the conceptual model for this species also demonstrated how new information from different sites could be integrated into the model when additional experts are included.

Similar minor updates were made to the conceptual model for the large bent-winged-bat, where uncertainty surrounding windfarms was removed based on new information that these would pose a serious threat to the species. For Caley’s grevillea, lack of genetic diversity was now considered as a threat, and translocation was added as a potential management intervention. In data-poor projects, such as Mahoney’s toadlet, reviewing the conceptual model allowed participants to update their assumptions based on new knowledge (e.g., the impact of *Gambusia* on tadpoles and chytrid fungus on adults) and identify new priorities (e.g., connectivity between core and peripheral sites). Updating the conceptual model provided a means to quantify the knowledge gained from extensive research on this species and prioritise intervention objectives such as maintaining habitat connectivity and controlling cats. The full list of updates to the conceptual models is given in Supplementary Table S1.

For the two species where the observed outcome was worse than expected, the conceptual models were used to explore the potential reasons for the discrepancy. In the case of the long-nosed potoroo at the Barrington Tops site (Fig. 3), the conceptual model was used to explore reasons why a reduction in foxes may not have resulted in an increase in the potoroo population. Potential reasons included a reduction in foxes leading to an increase in cats, the site already being at carrying capacity, or it may be that the choice of occupancy as an indicator was not sensitive to an increase in abundance. Similarly, the conceptual model for the tall rusthood orchid was instrumental in showing why the planned management actions hadn’t worked by highlighting the unforeseen impact of herbivory from white-winged choughs (*Corcorax melanorhamphos*) and the lack of winter grazing in the caged areas (increased competition from other plants and herbivory from slugs).

**Fig. 3.**
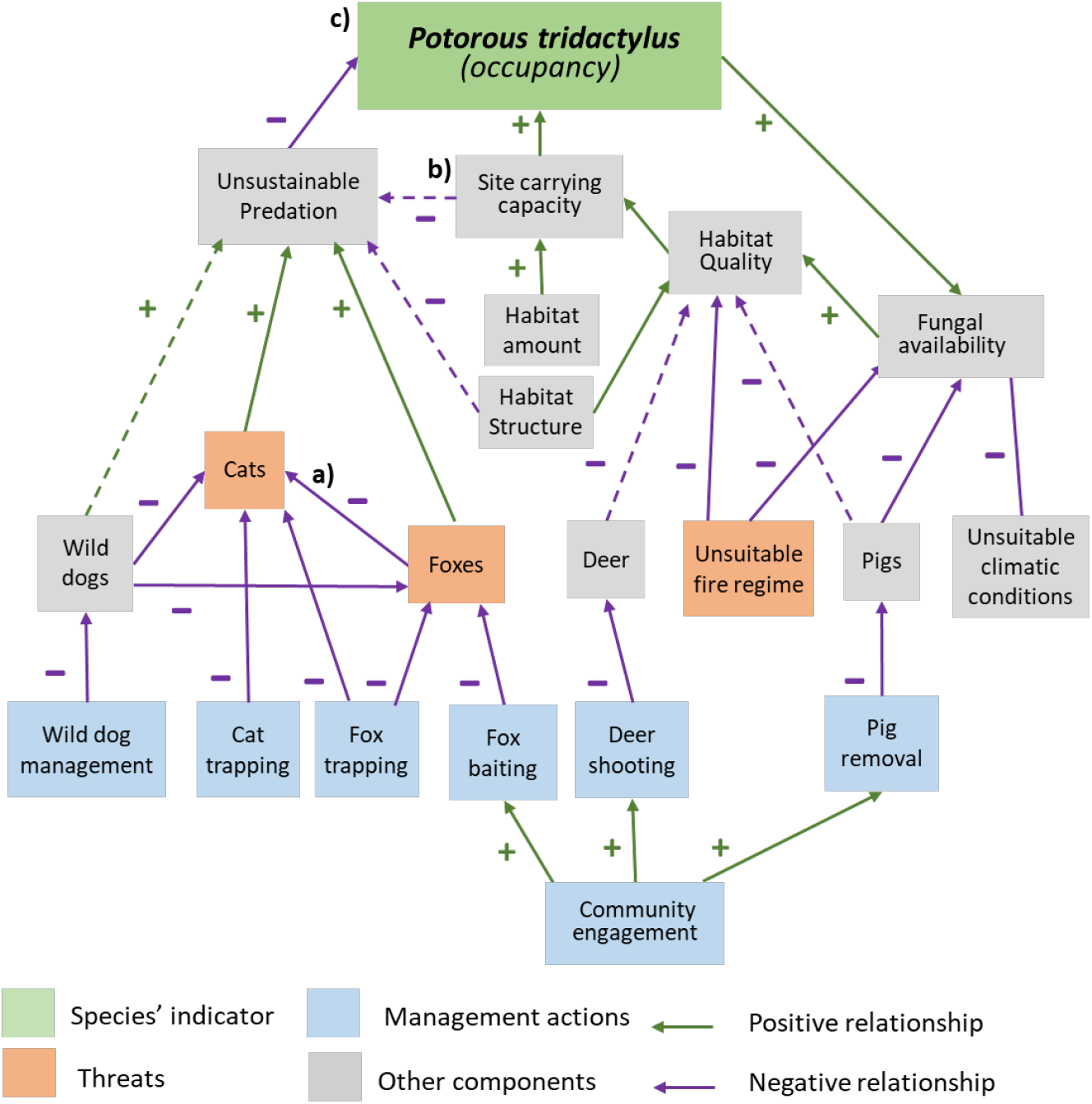
Updated conceptual model for the long-nosed potoroo Saving Our Species (SoS) management plan which shows potential reasons why a decrease in foxes may not result in increased potoroos because of a) a possible increase in cats, b) site carrying capacity c) sensitivity of selected indicator

### Practicality of the approach

Of the 13 participants in the review (six project coordinators and seven additional experts), five had been directly involved in the original development of the guidelines, including creating the conceptual models and response to management curves for their project. These experts consistently reported that the structured approach had influenced their management decisions over the five years from 2019 to 2023. However, they emphasised the importance of periodic review, noting that models and estimates need to be updated as new information emerges and as site-specific conditions change.

The inclusion of eight participants who were not involved in the original case study design, highlighted the utility of the conceptual models as tools for communicating and synthesising existing knowledge. However, they noted that the usefulness of these models depended on having well-defined project scope to ensure meaningful interpretation. For example, experts identified that the broad-headed snake has different behaviours at different sites, depending on the distance of the trees to the basking rocks. This difference impacts the which monitoring approaches are feasible.

In the two case studies where multiple experts participated in the review, conceptual models were considered particularly valuable in facilitating collaboration, especially when there were different perspectives. Experts described how visualising the logic using the models helped clarify differing interpretations of key relationships between species, threats and management actions. This was also true for multi-species projects like Floyd’s grass and the black grass-dart butterfly, a conceptual model provided a structured way to document the dependent relationships, helping ensure that management actions benefit both species rather than treating them separately.

For the two newly developed case studies that applied the guidelines to multi-species management (Supplementary material S4), participants described the five-step process as intuitive and easy to follow. Projector coordinators for these two case studies identified the inclusion of worked examples and the structured, stepwise approach to estimating management as key strengths.However, they also noted that constructing a narrative for the conceptual model was one of the more challenging aspects of the process, suggesting additional guidance is needed.

## Discussion

This review of eight threatened species case studies provides rare empirical insight into the utility of expert-elicited data in adaptive conservation planning. By revisiting structured conceptual models and predicted response to management estimates five years post-implementation, we were able to assess not only the alignment between predicted and observed outcomes, but also the practical value of the process in real-world conservation settings. The following discussion explores the insights gained, limitations encountered, and implications for strengthening the use of structured expert knowledge in dynamic, resource-constrained environments.

### Adaptive management insights

The evaluation of previous elicited values and assumptions demonstrated how integrating expert knowledge with monitoring data supports adaptive conservation planning for threatened species and ecosystems. Conceptual models, when developed early and reviewed at ecologically relevant intervals, provided a structured and transparent approach to synthesising available knowledge. Experts found that this process helped clarify assumptions and refine management responses over time. Additionally, retrospective use of models proved valuable, particularly to inform improved adaptive strategies. The ability of experts—both those initially involved and newly engaged—to update models quickly through brief meetings and email discussions suggests that the conceptual model approach is scalable and facilitates rapid integration of new information. This finding is particularly relevant for conservation programs, like Saving our Species, that require iterative adjustments in response to evolving threats or new ecological insights.

### Conceptual model strengths and decision making

The use of expert-predicted response curves was valuable regardless of whether observed outcomes aligned more closely with predicted management or counterfactual scenarios. While confirmation bias remained a risk, particularly in the absence of true control sites, expert predictions provided reference points that helped determine whether conservation actions were achieving their intended outcomes. Future work integrating formal decision triggers (Cook et al. 2016, de Bie et al. 2018) would enhance structured decision making and provide more objective thresholds for adaptive responses. In some scenarios such as the long-nosed potoroo and tall rusthood orchid, the observed outcomes did not align with the expert predictions, instead resembling unmanaged counterfactual scenarios. However, rather than dismissing these models as flawed, experts were able to systematically explore possible explanations, including unanticipated threats, incorrect assumptions, or unforeseen species responses. This highlighted a key strength of the conceptual modelling approach, as it provides a structured framework for generating and testing new hypotheses when unexpected outcomes occur.

A key insight from these case studies was the importance of clearly defining project scope and incorporating site-specific information into conceptual models. Experts noted that models were most effective when grounded in the ecological and operational realities of each site, which helped ensure that multi-species or multi-site projects remained interpretable and actionable. While in some instances a separate model for each site may be warranted, in most cases only minor differences exist such as the presence or absence of pest predators or variation in human disturbance. A practical approach is to define a generalised model that includes all key relationships and annotate where differences occur that would affect management. For example, in the long-nosed potoroo case study (Supplementary material S1), the general model was sufficient but required site-specific annotations to account for variation in predator pressure and human disturbance. This approach proved especially useful when observed outcomes diverged from predictions, allowing experts to explore site-based factors.

### Challenges in indicator selection and monitoring

Selecting appropriate monitoring indicators requires balancing ecological relevance, feasibility, and sensitivity to management interventions (Lindenmayer, Woinarski et al. 2020, Mayfield et al. 2022, Rose et al. 2023). Participants in this study noted that occupancy-based measures were often more practical than abundance estimates but conceded that there were trade-offs. While occupancy (based on cameras or traps) remains the most practical indicator for measuring many species, the limitations at a site level need to be factored into interpretation of monitoring results, i.e. is it sufficiently sensitive to pick up changes in population abundance that result from management interventions. In cases like the broad-headed snake, indicator selection was constrained by site-specific monitoring limitations, where abundance estimates were feasible at one location but not others. Indicator complexity is also a consideration, for example in this study only naïve occupancy was available for the long-nosed potoroo as resources were not yet available to construct an occupancy model from the data. In some instances, particularly for cryptic or mobile species, occupancy remains both the most practical and ecological suitable indicator (Mayfield, Bird et al. 2022). However, careful consideration is needed when shifting between abundance and occupancy metrics to ensure trends can be tracked through the transition period. Future conservation programs could benefit from testing both metrics in parallel where resources allow, focusing on transitions between monitoring approaches.

### Collaborative model development and stakeholder engagement

Collaborative model development proved critical for aligning project decisions with diverse ecological realities (Rose, Hemming et al. 2023). Engaging a broad range of experts helped ensure that key threats, dependencies, and site-specific variations were accounted for (Martin, Burgman et al. 2012). This was evident in the broad-headed snake project, where early models focused on only one key site until additional expert input revealed site-specific habitat influences on species dynamics. This supports the importance of structured, iterative, stakeholder engagement in adaptive conservation planning (Margoluis et al. 2009, Rose, Hemming et al. 2023). Early conceptual models should be viewed as living documents, updated as new evidence emerges or as site conditions change (Margoluis, Stem et al. 2009). Defining the project scope from the outset is particularly important for multi-species initiatives, where broader interactions can complicate interpretation and decision-making (Runge et al. 2011, Lindenmayer, Woinarski et al. 2020).

### Implications for conservation planning and future directions

This study provides a practical example of using structured expert judgment to inform adaptive conservation management. While the approach has proven valuable across a range of taxa and both single and multi-species conservation plans, several limitations should be considered. The accuracy of predicted outcomes depends on how well assumptions align with ecological realities, and confirmation bias can influence expert estimates, particularly in data poor contexts. To mitigate bias, models should be developed with diverse expert representation and assumptions should be clearly documented to ensure transparency. By systematically testing and updating these models over time, conservation practitioners can improve alignment between predictions and real-world outcomes, ultimately enhancing species recovery efforts and long-term conservation success.

Insights from this study reinforce the value of structured decision-making in conservation. Conceptual models, combined with expert elicitation, helped to prioritise uncertainties, ensuring that management decisions focus on the factors with greatest influence on conservation outcomes. By systematically mapping species interactions, threats, and management responses, these models enable practitioners to make informed predictions about conservation interventions, identify critical knowledge gaps requiring further research, and refine management strategies iteratively based on new evidence. Beyond structuring knowledge, conceptual models translated ecological complexity into clear, actionable steps. By breaking down intricate relationships into manageable components, they support conservation teams to identify urgent threats, optimal interventions, and the logical sequence of actions needed to achieve conservation goals. This approach fosters transparency, collaboration, and adaptability, ensuring that conservation planning remains responsive to emerging challenges.

## Supporting information

S1 - Updated case studies

S2 Table - Conceptual diagram updates

S3 Table - Observed v predicted vaules

S4 - Multi-species case studies

## Acknowledgements

We gratefully acknowledge the contributors to the original case studies and guidelines that were reviewed in this study, as well as the Saving our Species staff and other individuals who have worked to management these species over the last five years. Simon Clulow is acknowledged for early contributions to the U. mahonyi case study.

## Supplementary Files

Supplementary S1. Updates to original case studies

Supplementary S2 Table. Key changes to conceptual diagrams for each of the six review case studies

Supplementary S3 Table. Values for observes and predicted species indicators

Supplementary S4. Additional Case studies

## References

Adams-Hosking, C., M. F. McBride, G. Baxter, M. Burgman, D. de Villiers, R. Kavanagh, I. Lawler, D. Lunney, A. Melzer, P. Menkhorst, R. Molsher, B. D. Moore, D. Phalen, J. R. Rhodes, C. Todd, D. Whisson and C. A. McAlpine (2016). “Use of expert knowledge to elicit population trends for the koala (Phascolarctos cinereus).” Diversity and Distributions 22(3): 249–262.

Brazill-Boast, J., M. Williams, B. Rickwood, T. Partridge, G. Bywater, B. Cumbo, I. Shannon, W. J. M. Probert, J. Ravallion, H. Possingham and R. F. Maloney (2018). “A large-scale application of project prioritization to threatened species investment by a government agency.” PLOS ONE 13(8): e0201413.

Burgman, M. A. (2016). Trusting judgements how to get the best out of experts. Cambridge, UK, Cambridge University Press.

Camus, E. B., J. R. Rhodes, C. A. McAlpine, D. Lunney, J. Callaghan, R. Goldingay, A. Brace, M. Hall, S. B. Hetherington, M. Hopkins, M. J. Druzdzel and H. J. Mayfield (2022). “Using expert elicitation to identify effective combinations of management actions for koala conservation in different regional landscapes.” Wildlife Research: -.

Cook, C. N., K. de Bie, D. A. Keith and P. F. E. Addison (2016). “Decision triggers are a critical part of evidence-based conservation.” Biological Conservation 195(Supplement C): 46–51.

de Bie, K., P. F. E. Addison and C. N. Cook (2018). “Integrating decision triggers into conservation management practice.” Journal of Applied Ecology 55(2): 494–502.

Díaz, S., J. Settele, E. S. Brondízio, H. T. Ngo, J. Agard, A. Arneth, P. Balvanera, K. A. Brauman, S. H. M. Butchart, K. M. A. Chan, L. A. Garibaldi, K. Ichii, J. Liu, S. M. Subramanian, G. F. Midgley, P. Miloslavich, Z. Molnár, D. Obura, A. Pfaff, S. Polasky, A. Purvis, J. Razzaque, B. Reyers, R. R. Chowdhury, Y.-J. Shin, I. Visseren-Hamakers, K. J. Willis and C. N. Zayas (2019). “Pervasive human-driven decline of life on Earth points to the need for transformative change.” Science 366(6471): eaax3100.

Gregory, R. (2012). Structured decision making: a practical guide to environmental management choices. Hoboken N.J., Wiley-Blackwell.

Gumbs, R., O. Scott, R. Bates, M. Böhm, F. Forest, C. L. Gray, M. Hoffmann, D. Kane, C. Low, W. D. Pearse, S. Pipins, B. Tapley, S. T. Turvey, W. Jetz, N. R. Owen and J. Rosindell (2024). “Global conservation status of the jawed vertebrate Tree of Life.” Nature Communications 15(1): 1101.

Hemming, V., M. A. Burgman, A. M. Hanea, M. F. McBride and B. C. Wintle (2018). “A practical guide to structured expert elicitation using the IDEA protocol.” Methods in Ecology and Evolution 9(1): 169–180.

Hemming, V., A. E. Camaclang, M. S. Adams, M. Burgman, K. Carbeck, J. Carwardine, I. Chadès, L. Chalifour, S. J. Converse, L. N. K. Davidson, G. E. Garrard, R. Finn, J. R. Fleri, J. Huard, H. J. Mayfield, E. M. Madden, I. Naujokaitis-Lewis, H. P. Possingham, L. Rumpff, M. C. Runge, D. Stewart, V. J. D. Tulloch, T. Walshe and T. G. Martin (2022). “An introduction to decision science for conservation.” Conservation Biology 36(1): e13868.

Kingsford, R. T., H. C. Biggs and S. R. Pollard (2011). “Strategic Adaptive Management in freshwater protected areas and their rivers.” Biological Conservation 144(4): 1194–1203.

Kleitou, P., F. Crocetta, S. Giakoumi, I. Giovos, J. M. Hall-Spencer, S. Kalogirou, D. Kletou, D. K. Moutopoulos and S. Rees (2021). “Fishery reforms for the management of non-indigenous species.” Journal of Environmental Management 280: 111690.

Lanzas, M., N. Pou, G. Bota, M. Pla, D. Villero, L. Brotons, P. Sainz de la Maza, J. Bach, S. Pont, M. Anton, S. Herrando and V. Hermoso (2024). “Detecting management gaps for biodiversity conservation: An integrated assessment.” Journal of Environmental Management 354: 120247.

Lindenmayer, D., J. Woinarski, S. Legge, D. Southwell, T. Lavery, N. Robinson, B. Scheele and B. Wintle (2020). “A checklist of attributes for effective monitoring of threatened species and threatened ecosystems.” Journal of Environmental Management 262: 110312.

Margoluis, R., Salafsky, N (1998). Measures of success: designing, managing, and monitoring conservation and development projects, Island Press.

Margoluis, R., C. Stem, N. Salafsky and M. Brown (2009). “Using conceptual models as a planning and evaluation tool in conservation.” Evaluation and Program Planning 32(2): 138–147.

Martin, T. G., M. A. Burgman, F. Fidler, P. M. Kuhnert, S. Low-Choy, M. McBride and K. Mengersen (2012). “Eliciting Expert Knowledge in Conservation Science.” Conservation Biology 26(1): 29–38.

Mayfield, H., J. Rhodes, M. Evans and M. Maron (2019). Saving our Species Guidelines for Estimating and Evaluating Species’ Response to Management. Report to Department of Planning, Industry and Environment. Sydney, Australia.

Mayfield, H. J., J. Bird, M. Cox, G. Dutson, T. Eyre, K. Raiter, J. Ringma and M. Maron (2022). “Guidelines for selecting an appropriate currency in biodiversity offset transactions.” Journal of Environmental Management 322: 116060.

Mayfield, H. J., J. Brazill-Boast, E. Gorrod, M. C. Evans, T. Auld, J. R. Rhodes and M. Maron (2020). “Estimating species response to management using an integrated process: A case study from New South Wales, Australia.” Conservation Science and Practice 2(11): e269.

Mayfield, H. J., R. Eberhard, C. Baker, U. Baresi, M. Bode, A. Coggan, A. J. Dean, F. Deane, E. Hamman, D. Jarvis, B. Loechel, B. M. Taylor, L. Stevens, K. Vella and K. J. Helmstedt (2023). “Designing an expert-led Bayesian network to understand interactions between policy instruments for adoption of eco-friendly farming practices.” Environmental Science & Policy 141: 11–22.

Nichols, J. D. and B. K. Williams (2006). “Monitoring for conservation.” Trends in Ecology & Evolution 21(12): 668–673.

Rose, L. E., V. Hemming, A. M. Hanea, B. A. Wintle and Y. E. Chee (2023). “Linking species distribution models with structured expert elicitation for predicting management effectiveness.” Conservation Science and Practice 5(12): e13038.

Runge, M. C., S. J. Converse and J. E. Lyons (2011). “Which uncertainty? Using expert elicitation and expected value of information to design an adaptive program.” Biological Conservation 144(4): 1214–1223.

Schwartz, M. W., K. Deiner, T. Forrester, P. Grof-Tisza, M. J. Muir, M. J. Santos, L. E. Souza, M. L. Wilkerson and M. Zylberberg (2012). “Perspectives on the Open Standards for the Practice of Conservation.” Biological Conservation 155: 169–177.

Stenhouse, A., T. Perry, F. Grützner, P. Rismiller, L. P. Koh and M. Lewis (2022). “COVID restrictions impact wildlife monitoring in Australia.” Biological Conservation 267: 109470.

Tucker, M. A., A. M. Schipper, T. S. F. Adams, N. Attias, T. Avgar, N. L. Babic, K. J. Barker, G. Bastille-Rousseau, D. M. Behr, J. L. Belant, D. E. Beyer, N. Blaum, J. D. Blount, D. Bockmühl, R. L. Pires Boulhosa, M. B. Brown, B. Buuveibaatar, et al. (2023). “Behavioral responses of terrestrial mammals to COVID-19 lockdowns.” Science 380(6649): 1059–1064.

van Eeden, L. M., Nimmo, D., Mahony, M., Herman, K., Ehmke, G., Driessen, J., O’Connor, J., Bino, G., Taylor, M. and Dickman, C. R. (2020). Impacts of the Unprecedented 2019-2020 Bushfires on Australian Animals. Sydney, WWF-Australia.

Walters, C. J. and C. S. Holling (1990). “Large-Scale Management Experiments and Learning by Doing.” Ecology 71(6): 2060–2068.

Ward, M., A. I. T. Tulloch, J. Q. Radford, B. A. Williams, A. E. Reside, S. L. Macdonald, H. J. Mayfield, M. Maron, H. P. Possingham, S. J. Vine, J. L. O’Connor, E. J. Massingham, A. C. Greenville, J. C. Z. Woinarski, S. T. Garnett, M. Lintermans, B. C. Scheele, J. Carwardine, D. G. Nimmo, D. B. Lindenmayer, R. M. Kooyman, J. S. Simmonds, L. J. Sonter and J. E. M. Watson (2020). “Impact of 2019–2020 mega-fires on Australian fauna habitat.” Nature Ecology & Evolution.

