## Supplementary material for "Evaluating a structured expert elicitation approach for adaptive conservation: Lessons from five years in practice": S1 - Updated case studies

### Supplementary S1: Updated case studies

**Content reproduced from:**

***Conceptual models for conservation planning: Evaluation of the SoS modelling approach and guidelines Saving our Species***

Environment and Heritage
Department of Climate Change, Energy, the Environment and Water
Locked Bag 5022, Parramatta NSW 2124.

2024

#### Contents

1 Potorous tridactylus (Long-nosed Potoroo) 2

1.1 Summary 2

1.2 Updates to conceptual model 2

1.3 Tracking against indicators 4

2 Miniopterus orianae oceanensis (Large bent-winged bat) 7

2.1 Summary 7

2.2 Updates to conceptual model 7

2.3 Tracking against indicators 9

3 *Uperoleia mahonyi* (Mahoney's toadlet) 13

3.1 Summary 13

3.2 Updates to the conceptual model 13

3.3 Tracking against modelled indicators 15

3.4 Some Key updates on our understanding of Uperoleia mahonyi 15

4 Pterostylis chaetophora (Tall rusthood) 17

4.1 Summary 17

4.2 Updates to conceptual model 17

4.3 Tracking against modelled indicators 19

5 *Grevillea caleyi* (Caley's grevillea) 23

5.1 Summary 23

5.2 Updates to the conceptual model 23

5.3 Tracking against modelled indicators 25

6 Hoplocephalus bungaroides (Broad-headed snake) 28

6.1 Summary 28

6.2 Updates to the conceptual model 28

6.3 Tracking against modelled indicators 30

### 1 Potorous tridactylus (Long-nosed Potoroo)

Contributors: Trent Forge[1]

**Role/Affiliations:** [1] Assistant Project Officer, Blue Mountains Branch, New South Wales National Parks and Wildlife Service

#### *1.1 Summary*

Management and monitoring for the long-nosed potoroo have been carried out as planned at all sites. An undesirable fire occurred at one site, which has impacted potoroo occupancy.

Review of the case study has split one site into two (Mt Royal and Barrington tops) in line with management and monitoring. The link between deer/pigs and long-nosed potoroo populations is now considered unlikely. While both feral species are found at most sites, their numbers are insufficient to effect potoroos habitat quality. Of note is the Barrington Tops site, where fox numbers declined significantly, but without the expected associated increase in potoroo occupancy. Several possible reasons have been suggested for this. First, the decrease in foxes may have led to an increase in cats, as per the conceptual model. Second, if the site was already at carrying capacity, then there may not be scope for a population increase in potoroos, in which case maintaining habitat quality becomes a priority. Third, it is possible that there has been an increase in potoroos that was not picked up by the monitoring, either because of the location of the camera traps or the indicator being used (occupancy). This risk was identified in the original case study which commented that potoroos are more likely to increase density around known localities than to disperse more widely. Other options could be that the high suitability of habitat structure limits the impact of predation, or that this represents a reversal of a declining trend.

#### *1.2 Updates to conceptual model*

The amount of new information on this species has led to significant updates to the conceptual model (Fig. 1)

- The link between deer/pigs and long-nosed potoroo populations is now considered unlikely, but the tentative link has been left in the model to indicate that this could change if populations of feral animals increase.
- Habitat quality and structure have been added to the model, albeit with some uncertainty indicated. There is now some possibility that habitat characteristics could be limiting predation, however more information is needed before this can be confirmed.
- The importance of cats, wild dogs, foxes and pigs has been identified as differing at each site. This information has been included in the conceptual model as a table indicating the strength of the link for each site.

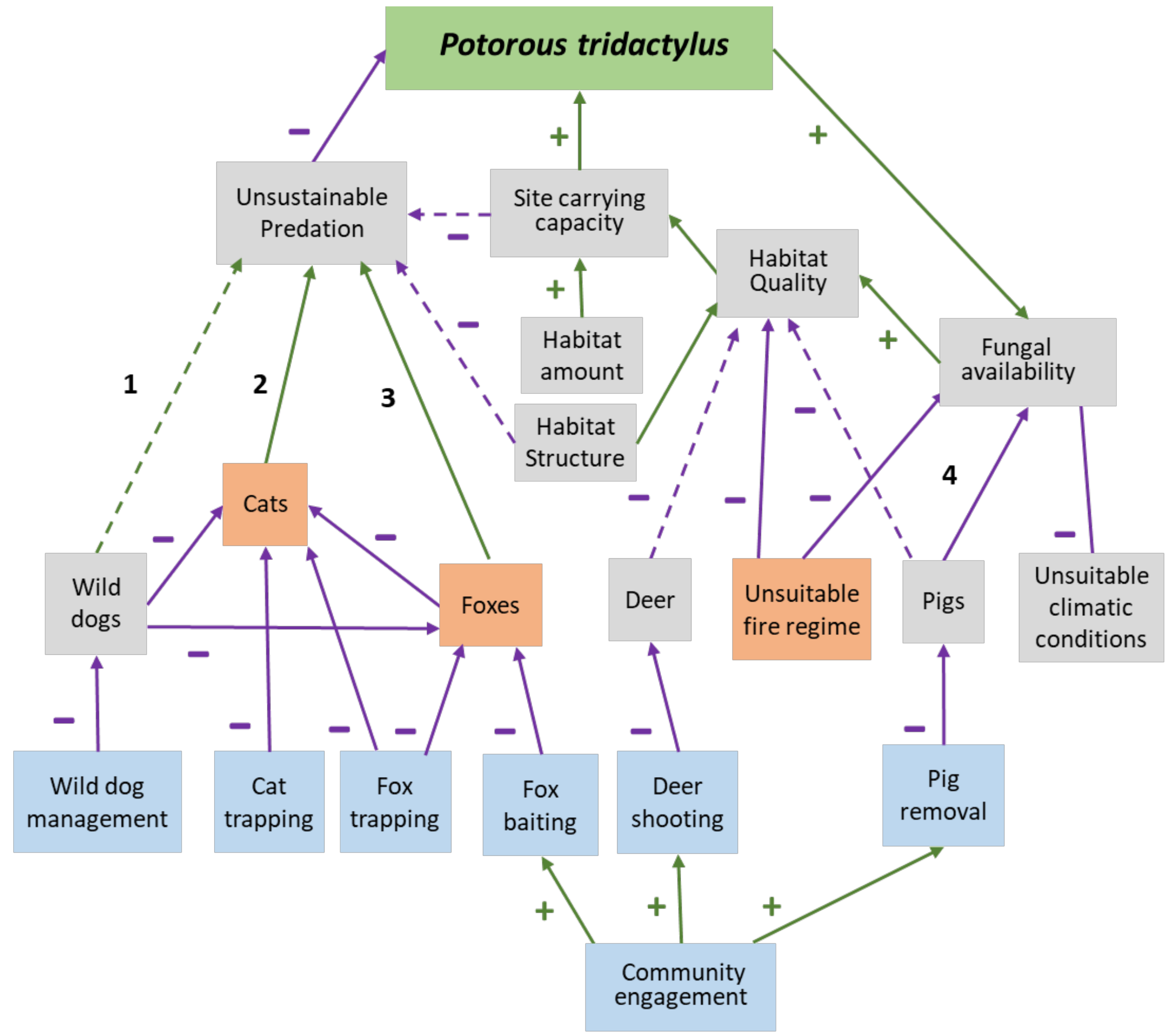


| Site | 1. Wild dogs | 2. Cats | 3. Foxes | 4. Pigs |
|---|---|---|---|---|
| Barrington tops | | | | |
| Budderoo/Barren grounds/Kangaroo Valley | | | | |
| South East | | | | |
| Mt Royal | | | | |
| Richmond Range | | | | |

**Fig. 1:** Updated conceptual model for the long-nosed potoroo

#### 1.3 Tracking against indicators

The indicators selected in 2018 were potoroo occupancy, fox occupancy, cat occupancy and habitat under suitable fire regime (Fig. 2)

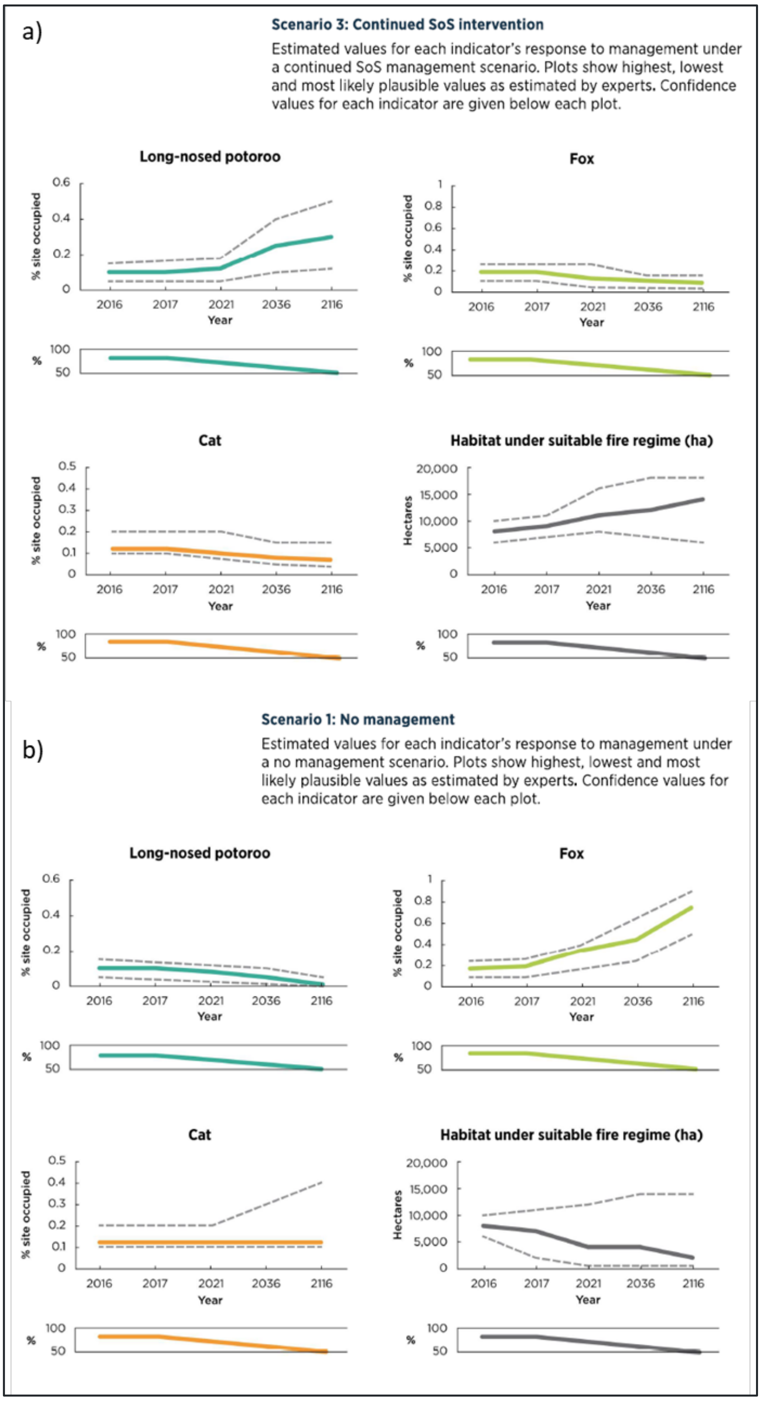


**Fig. 2: Original indicators and estimated response to management curves for the 2018 long-nosed potoroo case study for a) an SoS managed scenario and b) an unmanaged counterfactual.**

Occupancy rates for LNP at Barren Grounds are withing the range estimated by experts (Tables 1 & 2). At the Barrington Tops site, experts advised that fox numbers followed the expected trend and decreased with baiting (Table 3), however potoroo occupancy (not shown) did not increase as expected. Important to note although potoroo number is didn't decline, as predicted in the no management scenario, this could be for a number of reasons such as impact of habitat quality, or an increase in other predators such as cats. Comprehensive monitoring was carried out for the LNP including occupancy (captures per night and using camera traps), and occupancy for foxes and cats. Dingos were monitored sporadically at one site. Fire was not monitored but is considered in the notes for the other indicators.

**Table 1:** Naive occupancy data (camera traps) for the long-nosed potoroo. Data sourced from the SoS monitoring database

| Year | Occupancy - Camera traps | | | | |
|---|---|---|---|---|---|
| | Barren Grounds | Budderoo | Mount Royal - | Barrington Tops | South East Forests |
| 2016-2017 | 120/193 (74%) | - | - | - | - |
| 2017-2018 | 64% | - | - | - | - |
| 2018-2019 | 71% | - | 75% | - | 10% |
| 2019-2020 | 66% | - | - | - | 25% (Northern section)<br>10% (Southern section) |
| 2020-2021 | - | - | - | - | - |
| 2021-2022 | 65% | - | 50% | - | - |
| 2022-2023 | 55% | - | - | - | - |

**Table 2:** Naive occupancy data (captures) for the long-nosed potoroo. Data sourced from the SoS monitoring database

| Year | Occupancy - Captures per night | | | | |
|---|---|---|---|---|---|
| | Barren Grounds | Budderoo | Mount Royal | Barrington Tops | South East forests |
| 2016-2017 | 0.05, 0.01 | 0.04. 0.03 | - | - | - |
| 2017-2018 | 0.04 | 0.01 | 35% (all habitat), 70% (optimal habitat) | 25% (all habitat), 65% (optimal habitat) | - |
| 2018-2019 | 0.03 | 0.02 | - | - | - |
| 2019-2020 | - | - | - | - | - |
| 2020-2021 | - | - | - | - | - |
| 2021-2022 | - | - | - | - | |
| 2022-2023 | 0.03 | 0.02 | - | - | |

**Table 3:** Occupancy data for foxes at long-nosed potoroo management sites. Data sourced from the SoS monitoring database

| Year | Foxes | | | | |
|---|---|---|---|---|---|
| | Barren Grounds | Budderoo | Mount Royal - | Barrington Tops | South East Forests |
| 2016-2017 | 4% | 20% | - | - | - |
| 2017-2018 | 4% | 27% | 10% | 100% | - |
| 2018-2019 | 0% | 23% | 10% | - | 14% |
| 2019-2020 | 13% | 26% | - | - | 17% |
| 2020-2021 | - | - | <0.01% | 98% evenings | - |
| 2021-2022 | 8% | 20% | - | 0% (GT) | |
| 2022-2023 | 4% | 17% | - | - | |

**Table 4:** Occupancy data for cats at long-nosed potoroo management sites. Data sourced from the SoS monitoring database

| Year | Cats | | | | |
|---|---|---|---|---|---|
| | Barren Grounds | Budderoo | Mount Royal - | Barrington Tops | South East Forests |
| 2016-2017 | - | - | - | - | - |
| 2017-2018 | - | - | 100% | 80% | - |
| 2018-2019 | - | - | 100% | - | 4% |
| 2019-2020 | - | - | 40% of visits | | 8% |
| 2020-2021 | - | - | 48% of visits | | - |
| 2021-2022 | - | - | > 50% of visits | | |
| 2022-2023 | - | - | - | | |

**Table 5:** Occupancy data for dingoes at long-nosed potoroo management sites. Data sourced from the SoS monitoring database

| Year | Dingos | | | |
|---|---|---|---|---|
| | Barren Grounds | Budderoo | Mount Royal - | Barrington Tops |
| 2016-2017 | - | - | - | - |
| 2017-2018 | - | - | 100% | 80% |

### 2 Miniopterus orianae oceanensis (Large bent-winged bat)

Contributors: Doug Mills

**Role/Affiliations:** [1] Senior Project Officer Ecologist, New South Wales National Parks and Wildlife Service

#### 2.1 Summary

Management and monitoring for the large bent-winged bat (*Miniopterus* schreibersii *oceanensis*) was implemented as planned between 2018 and 2023. No additional threats have been identified, however the importance and certainty of several previously identified threats has changed. Specifically, there is now more evidence of the negative effect of windfarms on microbats, although no management interventions are known to mitigate this, and the effect is dependent on the distance between the roost and the turbines. The impact of cats on predation is now considered negligible, as very few cats are on site and they would not feasibly have capacity to take more than a couple of hundred bats out of a colony of tens of thousands. Resources for cat management and monitoring might be more efficiently used elsewhere. The risk of fungal pathogens, and their risk to this species remains uncertain.

#### 2.2 Updates to conceptual model

Additional evidence since 2018 has led to minor updates to the conceptual model (Fig. 3)

- The link from windfarms to bats has gone from uncertain to certain, and the impact of distance of turbines to roost has been included.
- Whether on not there are cats present is now shown as a dashed line to indicate a potentially negligible effect on predation

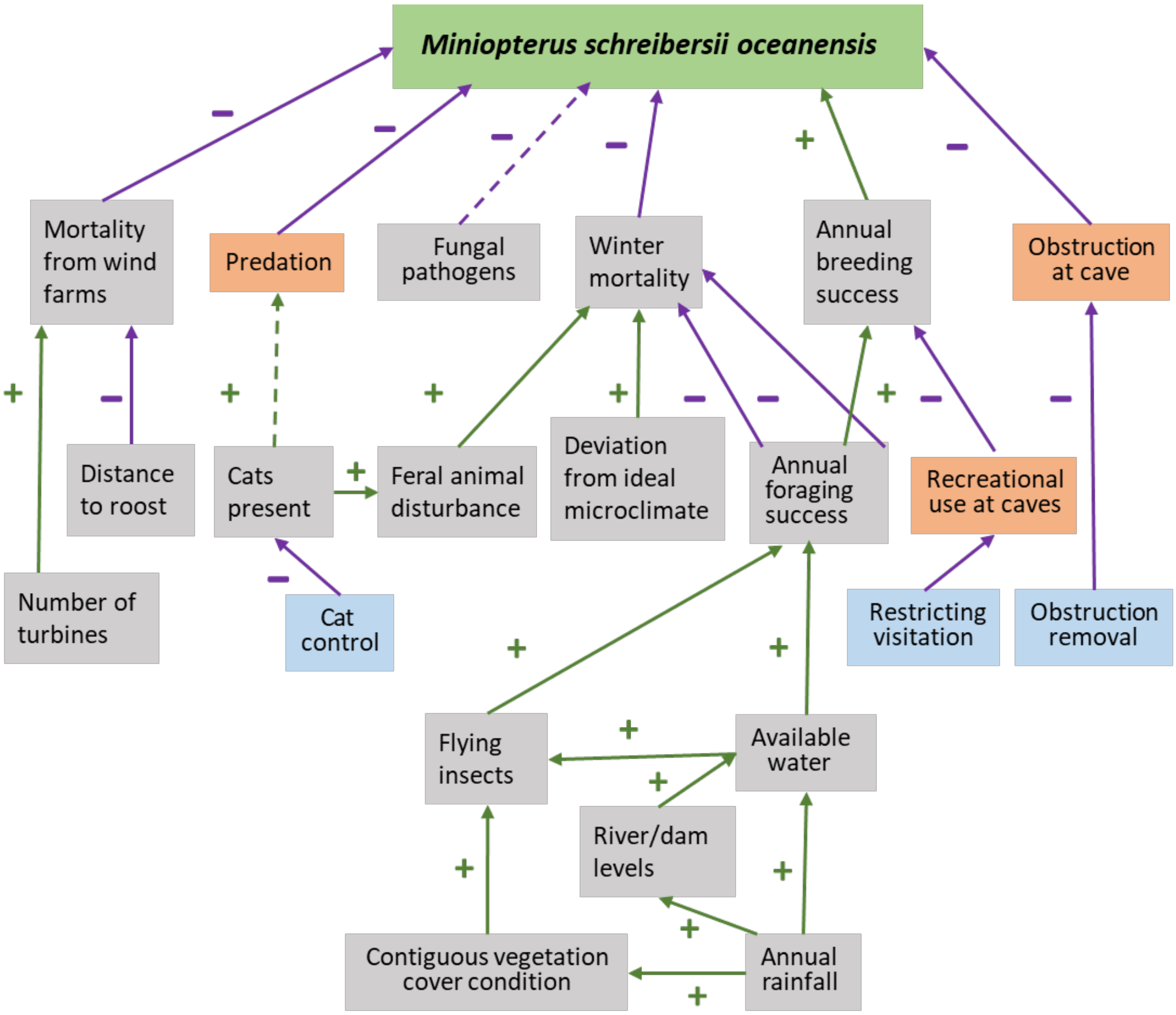


**Fig. 3:** Updated conceptual model for large bent-winged bat

#### 2.3 Tracking against indicators

The indicators selected in 2018 were number of bats, proportion of roost entrance covered, and nights per year with cats present (Fig. 4).

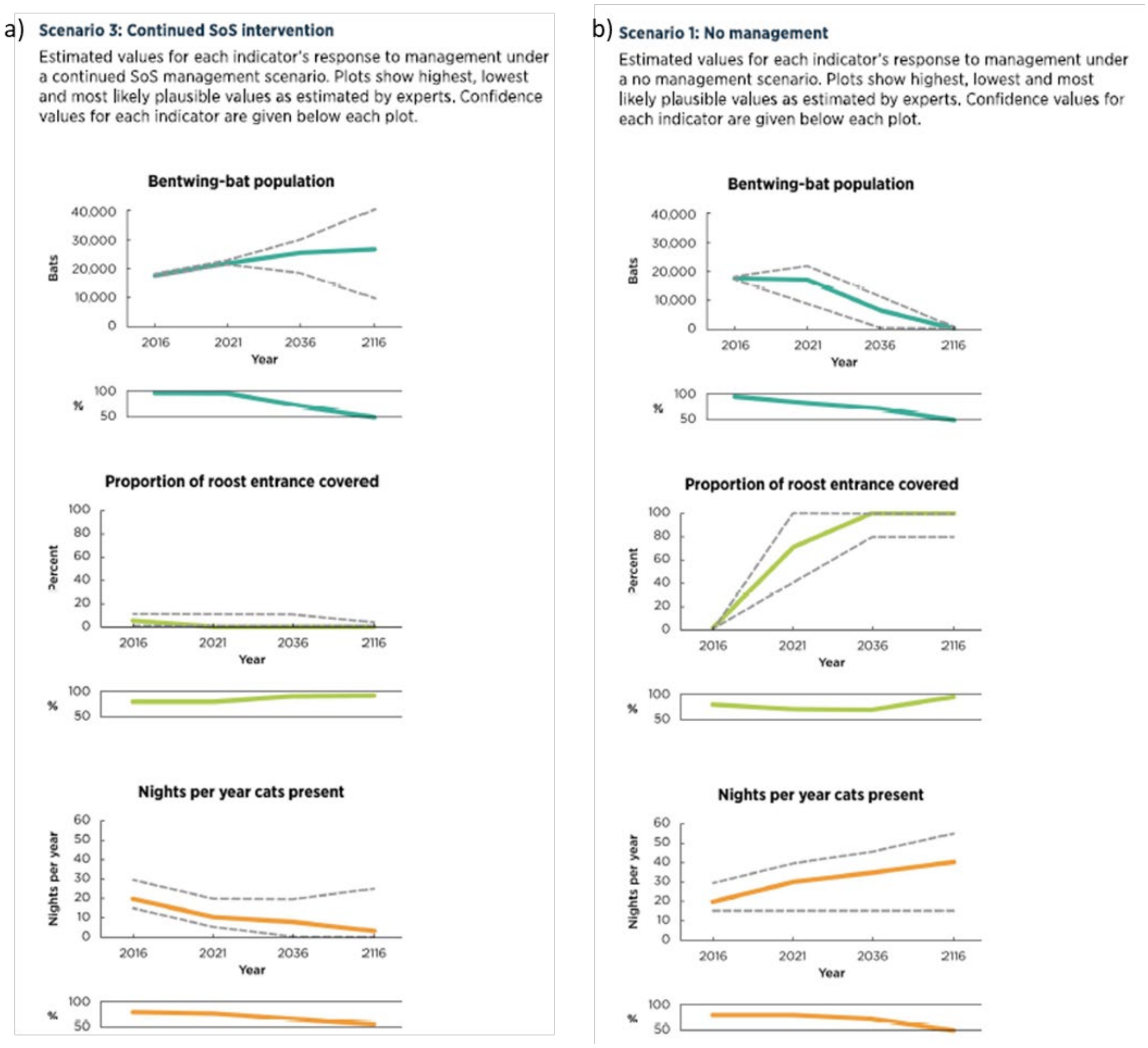


**Fig. 4:** Original indicators and estimated response to management curves for the 2018 large bent-winged bat case study for the a) Sos management scenario and b) No management scenario

The indictors that were monitored for the large bent-winged bat were abundance, cat activity, obstruction from vegetation and disturbance by people. Cat activity was measured as number of cats observed, rather than number of nights that cats were present, which allows for opportunistic sightings by rangers to be included. The Church Cave site (Fig. 4) was the basis for the original estimates, but all sites reflected an overall stable trend in population (Table 6)

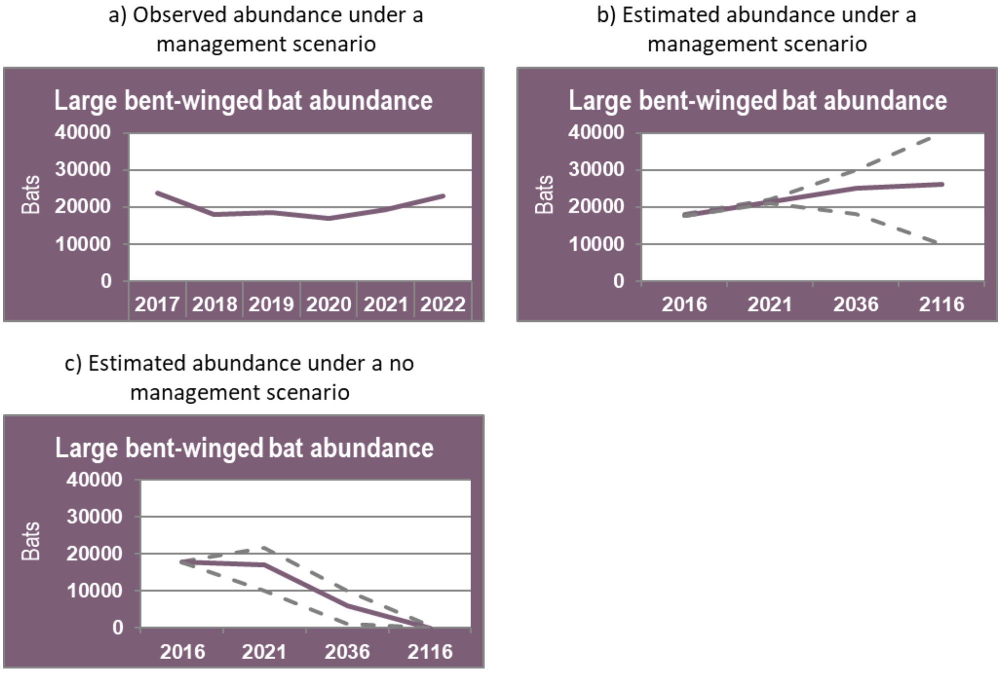


**Fig. 5:** Abundance values for the large bent-winged bat at the Church Cave site for a) observed values, b) estimated values for a managed scenario, and c) estimated values for an unmanaged scenario.

**Table 6:** Observed values for the large bent-winged bat population.

| **Year** | **Site** | | | | |
|---|---|---|---|---|---|
| | Church Cave | Dip Cave | Drum Cave | Kwiamble | Mount Kaputar |
| 2017-2018 | 23,700 ± 420 | - | - | - | - |
| 2018-2019 | 18,800 ± 570 | < 200 | 18,000 ± 2000 | - | 1437 |
| 2019-2020 | 18,350 ± 330 | < 200 | 20,000 ± 2000 | - | - |
| 2020-2021 | 16,900 ± 400 | < 200 | 16,000 ± 1500 | 114 | > 2000 |
| 2021-2022 | 19,200 | < 200 | - | 726 | > 2000 |
| 2022-2023 | 23,300 | - | - | 1095 | > 2000 |

Management actions have been successful in controlling cats, entrance encroachment and disturbance from people visiting the caves, and monitoring values (Tables 7-9) show a negligible impact from these threats. Observed values for all indicators are within the ranges of those predicted by experts for the management scenario.

**Table 7:** Observed values Vegetation encroachment of the cave entrance (post-management)

| Year | Vegetation encroachment of the cave entrance (%) | | | | |
|---|---|---|---|---|---|
| | Church Cave | Dip Cave | Drum Cave | Kwiamble | Mount Kaputar |
| 2017-2018 | 0% | - | - | - | - |
| 2018-2019 | < 5% | < 10 % | 0% | - | 0% |
| 2019-2020 | < 10% | - | 0% | 0% | 0% |
| 2020-2021 | 0% | - | 0% | 0% | 0% |
| 2021-2022 | 0% | - | - | 0% | 0% |
| 2022-2023 | 0% | - | - | < 5% | < 5% |

**Table 8:** Observed values for vegetation encroachment of the cave entrance (post-management)

| Year | Cats detected (number of cats) | | | | |
|---|---|---|---|---|---|
| | Church Cave | Dip Cave | Drum Cave | Kwiamble | Mount Kaputar |
| 2017-2018 | No cats detected | - | - | - | - |
| 2018-2019 | One cat detected | No cats detected | No cats detected | One cat detected | No cats detected |
| 2019-2020 | Cats detected on two nights | - | No cats detected | - | - |
| 2020-2021 | One - two cats detected on four nights. | - | No cats detected | - | - |
| 2021-2022 | One cat detected | - | - | No cats detected | - |
| 2022-2023 | No cats detected | - | - | No cats detected | No cats detected |

**Table 9:** Observed values for disturbance at caves from people

| Year | Disturbance at caves from people | | | | |
|---|---|---|---|---|---|
| | Church Cave | Dip Cave | Drum Cave | Kwiamble | Mount Kaputar |
| 2017-2018 | No disturbance | - | - | - | |
| 2018-2019 | No disturbance | No disturbance | No disturbance | Minimal disturbance | No disturbance |
| 2019-2020 | No disturbance | - | No disturbance | Visitation but no disturbance | - |
| 2020-2021 | No disturbance | - | No disturbance | Very high visitation | Minimal disturbance |
| 2021-2022 | No disturbance | - | - | 780 visitors over 3 month period | No disturbance |
| 2022-2023 | No disturbance | - | - | Threat remains present. | No disturbance |

### 3 *Uperoleia mahonyi* (Mahoney's toadlet)

Contributors: Luke Foster[1], Grant Webster[2]

**Role/Affiliations:** [1] Species Project Coordinator, Ecosystems and Threatened Species - Hunter Central Coast Branch, Saving our Species Program, NSW Department of Environment, Energy, Climate Change and Water and, [2] Species Expert, University of Newcastle

#### *3.1 Summary*

In 2018 Mahony's toadlet (*Uperoleia mahonyi*) was assigned to the Data Deficient management stream by the SoS program. Subsequently, and in collaboration with Dr Simon Clulow and Grant Webster, critical research actions were identified to improve the understanding of the species ecology, distribution and management requirements.

Since project implementation, survey and monitoring has found species populations are likely currently stable within the boom-and-bust cycle, but management is still required, particularly for the populations that occur outside of the reserve system and may be impacted by coastal development and sand mining, or have an extremely small extent of occurrence. SoS funded research has been crucial to address knowledge gaps for the conservation project, and provide the information needed to identify appropriate management actions and design a fit for purpose monitoring plan.

#### *3.2 Updates to the conceptual model*

The amount of new information on this species has led to significant updates to the conceptual model (Fig. 6), change include:

- Connectivity between core and peripheral sites is considered crucial for populations to react to seasonal environmental factors.
- Certain threats (coastal development, sand mining,) are not relevant for the population in a State protected area.
- Juveniles, and to some extent adults, appear to have a boom and bust cycle in the ephemeral ponds, but are evidently more stable in the core sites, although with noticeable fluctuations in response to environmental factors (mainly drought). In the last few years population has apparently increased as it comes out of a drought cycle.
- Threats differ for each population. The Myall Lakes population, for example, would probably be ok if left unmanaged.
- The species can persist in a modified landscape, and is tolerant to some disruption, as long as the hydrology is suitable. Habitat suitability is driven primarily by geology and soil, and if these are not suitable, other species will occupy the site.
- We don't know exactly why the sand is important. Possibly due to causing lower pH of soil and/or water, and resulting impacts on development of frog eggs/tadpoles, as is the case for other frog species.
- Water and soil chemistry are known to affect the presence, and survival rates, of eggs and tadpoles for frogs generally (although specific tolerances for *U. mahonyi* are presently unknown), and likely plays and important role for *U. mahonyi* as they occur in ponds or wetlands that are supplied through ground water, primarily on Quaternary sands.

- Water extraction is a possible threat at some sites, particularly during droughts when wetlands are naturally drier and water extraction may cause further water table drawn down. Pigs are no longer considered a threat.
- Gambusia are now considered a potential threat (previously uncertain), although they are largely found in permanent water bodies, and frogs will usually avoid ponds where there are present.
- Deer were discussed and considered not to be an issue.
- Chytrid is now considered a potential threat (previously uncertain), as it has been identified in some sub-populations.

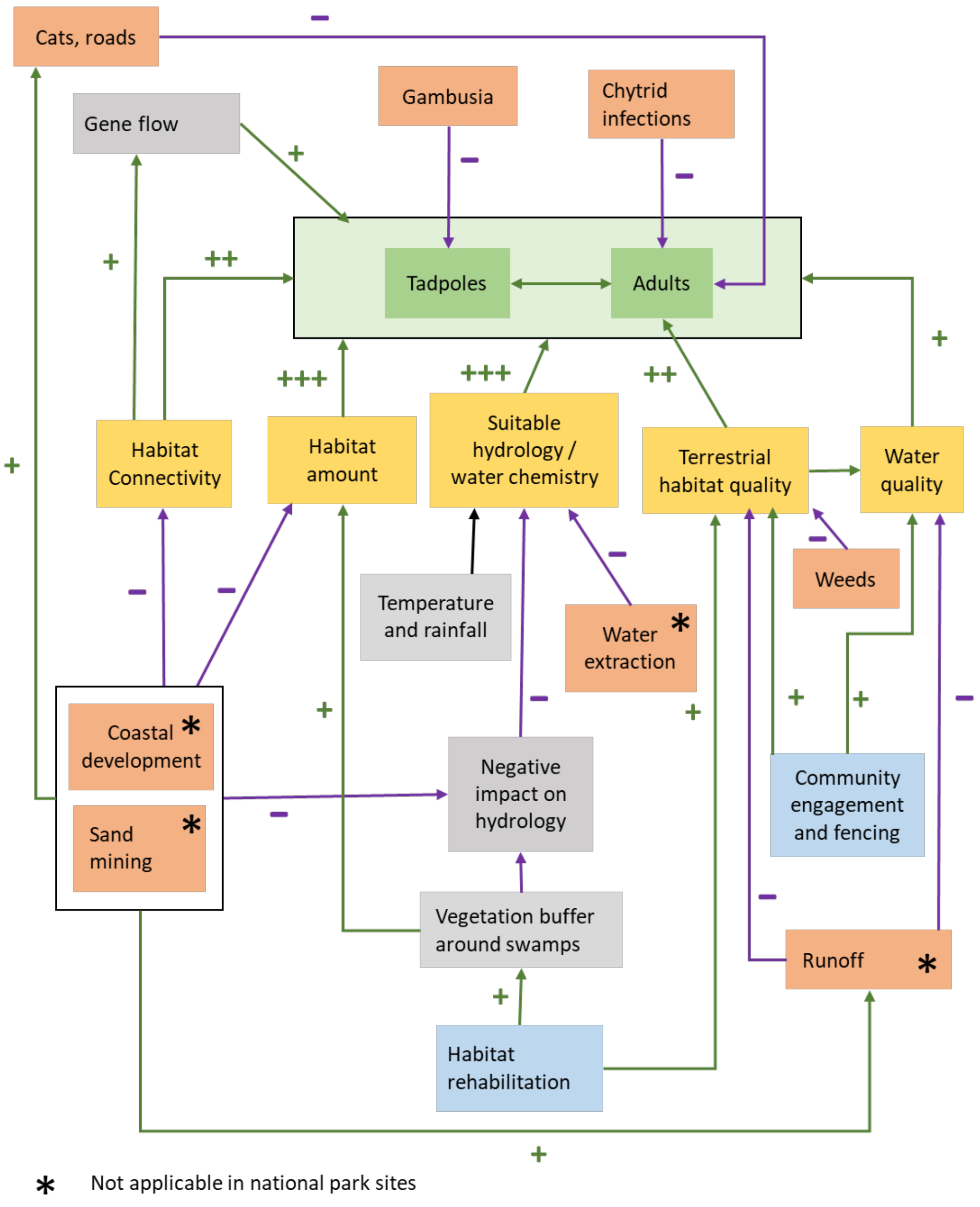


**Fig. 6:** Updated conceptual model for Mahoney's toadlet

#### 3.3 Tracking against modelled indicators

The original indicators selected for the Mahoney's toadlet are given in Table 10, along with the updates based on our improved understanding of the species.  A key consideration for monitoring for this species is the importance of peripheral sites. These are crucial for populations to expand, and contract based on environmental conditions. Absences of frogs at a peripheral site doesn't mean that the site doesn't requirement management or protection.

**Table 10:** Original indicators selected for Mahoney's toadlet including suggested updates based on our improved understanding of the species.

| Indicators | Trend | Update |
|---|---|---|
| Original (2018) | | |
| Frog population | Increased (due to increased survey efforts) | Based on our improved knowledge, this indicator has been changed to **number of calling males**, with caveats on correct environmental conditions for surveys. |
| Count of tadpoles | Not measured (intensive survey effort required) | This is important to confirm breeding activity and occupancy, however this direct measure is labour intensive and costly. Courser monitoring measures (e.g. eDNA) are likely similarly appropriate to indicate breeding activity and presence/absence. |
| Water quality score | Not measured (but may be in the near future) | It is likely that this will be physical measurements (e.g. pH, contaminants) rather than the more subjective water quality score defined in 2018. |
| Damage by pigs | No longer considered relevant and was not monitored. | To avoid unnoticed loss in a population, new estimates as provided for each of the four genetically distinct populations. |

#### 3.4 Some Key updates on our understanding of Uperoleia mahonyi

Improved understanding of the species general ecology

- *U. mahonyi* is associated with Quaternary geology, being coastal sediments (both Pleistocene and Holocene) and almost entirely sands (with some very minor clay and alluvium) – totalling 12 natural geologic units. Sands are frequently coastal (and ocean) deposited, reworked as aeolian dunes and subsequently eroded. Some occurrences are on modified (anthropogenic) geologies, amongst otherwise suitable Quaternary sands, indicating some tolerance to disturbed environments. The species has never been recorded from geologies that pre-date the Quaternary Period, even in close proximity to existing site occurrences, nor has it been recorded from any (natural) geology that does not constitute a sedimentary deposit.
- *U. mahonyi* is also associated with specific soil types that are associated with the sedimentary geologies, often sandy soils (both podzols and siliceous) and acid peats within wetland systems. It is known from only 6 natural soil types.

- Geology and soil explain *U. mahonyi* site occupancy; geology and climate explain U*. mahonyi* occurrence records; while geology, soil and climate explain the geographic range of *U. mahonyi*. Although *U. mahonyi* occurs in specific vegetation communities, vegetation type does not explain its distribution; rather vegetation and *U. mahonyi* are likely correlated due to collinearity driving shared preferences for geology, soil, and climate.
- Ponds and wetlands occupied as breeding sites are near-ubiquitously ephemeral or semi-permanent, artificial permanent ponds are also used infrequently. Breeding sites are apparently groundwater dependent and inundate as the water table rises following heavy rainfall events. The Quaternary sand systems that *U. mahonyi* inhabits are frequently underlaid by significant aquifers. Given the low-relief of the region *U. mahonyi* occupies, large areas of suitable wetlands exist during times of abundant rainfall.

Evaluating survey effort and improving detection probability

- Detection probability has been modelled, allowing for the provision of appropriate surveying advice practical to both policy makers and applied ecologists. Increased detectability is highly dependent on greater local abundance and breeding pond fullness.
- Two “activity seasons” exist: a breeding season; and a non-breeding season. The breeding season is extensive, ranging from July through to April and May. During the breeding season U. mahonyi forms loud choruses that are highly detectable; while choruses do not form in the non-breeding period. Chorus formation is strongly dependent on pond fullness, and choruses do not form when ponds are dry or close too. Given this, pond fullness is only important for increased detection during the breeding season, during the non-breeding season increased pond fullness does not result in higher detection probability.
- Increased pond fullness and heavy rainfall events explain increased calling activity, especially at ephemeral and semi-permanent sites. Regular calling occurs for as long as breeding ponds are full, or near full, continuously for consecutive months if conditions are suitable. At highly ephemeral sites breeding activity is more sporadic and infrequent. While calling at permanent sites is less variable and associated with rainfall.
- Surveying should be conducted during the breeding season, ideally when ponds are full, and should not be conducted if ponds are <60-70% full. At a site with average frog abundance, a minimum of two one-hour surveys are required to reach 95% confidence of a true absence, if conducted during the breeding season when the pond is at capacity. When ponds are dry, seven surveys during the breeding season would be required. Up to 15 surveys may be required at a site where the species is assumed to be rare.
- Surveying should not be conducted during the non-breeding season, as easily identifiable breeding choruses do not form and detections are restricted to visual observations or aural identification of irregular, non-advertisement, vocalisations, that may be territorial calls. These calls are not easily distinguished from sympatric myobatrachids and surveying during the non-breeding season creates considerable potential for site false negatives.

### 4 Pterostylis chaetophora (Tall rusthood)

Contributors: Paul Hillier

**Role/Affiliations:** [1] Species Project Coordinator, Ecosystems and Threatened Species - Hunter Central Coast Branch, Saving our Species Program, NSW Department of Environment, Energy, Climate Change and Water

#### *4.1 Summary*

In 2018, the critical threats identified to the tall rusthood orchid (*Pterostylis chaetophora*) at the Black Creek Saving our Species (SoS) priority management site were related to herbivory by native and feral mammals and competition from other native?invasive? vegetation, as well as some risk from trampling by xx. After five years of implementation, several substantial new threats have emerged which will be included in the model, and included in the project, the most unforeseeable of which is the detrimental behaviour of a family of white-winged choughs (*Corcorax melanorhampho*s). In 2019, the choughs began to eat the tubers of the plant, presumably in response to a food shortage during the drought. Additionally, a major development threat has emerged nearby, and while the species was not known at the site at the time of approval, and therefore not included in the offset arrangements, translocation of individuals may be practical if plants are found within the approved development area. The final major change to the model is to management actions. Due to lack of capacity to revisit the site within a season, cages at some sites were left out over winter, preventing winter "grazing" under the cages. Whilst the additional growth should not have impacted the viability of the orchid, the microclimate created in these localised patches attracted slugs, which preferentially ate orchid stems before they could flower. The management strategy has been amended to prioritise removing the cages each season to allow for winter grazing.

#### *4.2 Updates to conceptual model*

The change in the threats and pressures on this species has led to significant updates to the conceptual model (Fig.7) including:

- Grazing has been separated into growth-period (Aug-Jan) and dormant period (Feb-July) grazing. Grazing for February to July before the early budding phase removes competition from shrubs and weeds, as well as the dense groundcover that attracts slugs within the cages. Caging has been split into dormant period and growth-period to better define the management response.
- The new threat of nearby development has been added, having effects on habitat loss (although notably not at the site itself), direct mortality, trampling and increased summer and winter grazing macropods displaced by adjacent habitat loss.
- The threat from tuber-eating choughs has been added, which is more common in drier conditions, and can be managed to some extent with cages, however other more scalable, novel approaches are being sought.

Due to a population surviving an intense grass fire in a non-SoS site in 2019 we know that a single fire event is not necessarily detrimental to this species. At the Black Creek site, there was a Cultural Burn in May 2021. Again this was a single event, but observationally it did improve weeds for a time. Therefore, burning can be used as a tool to manage native and exotic groundcover competition.

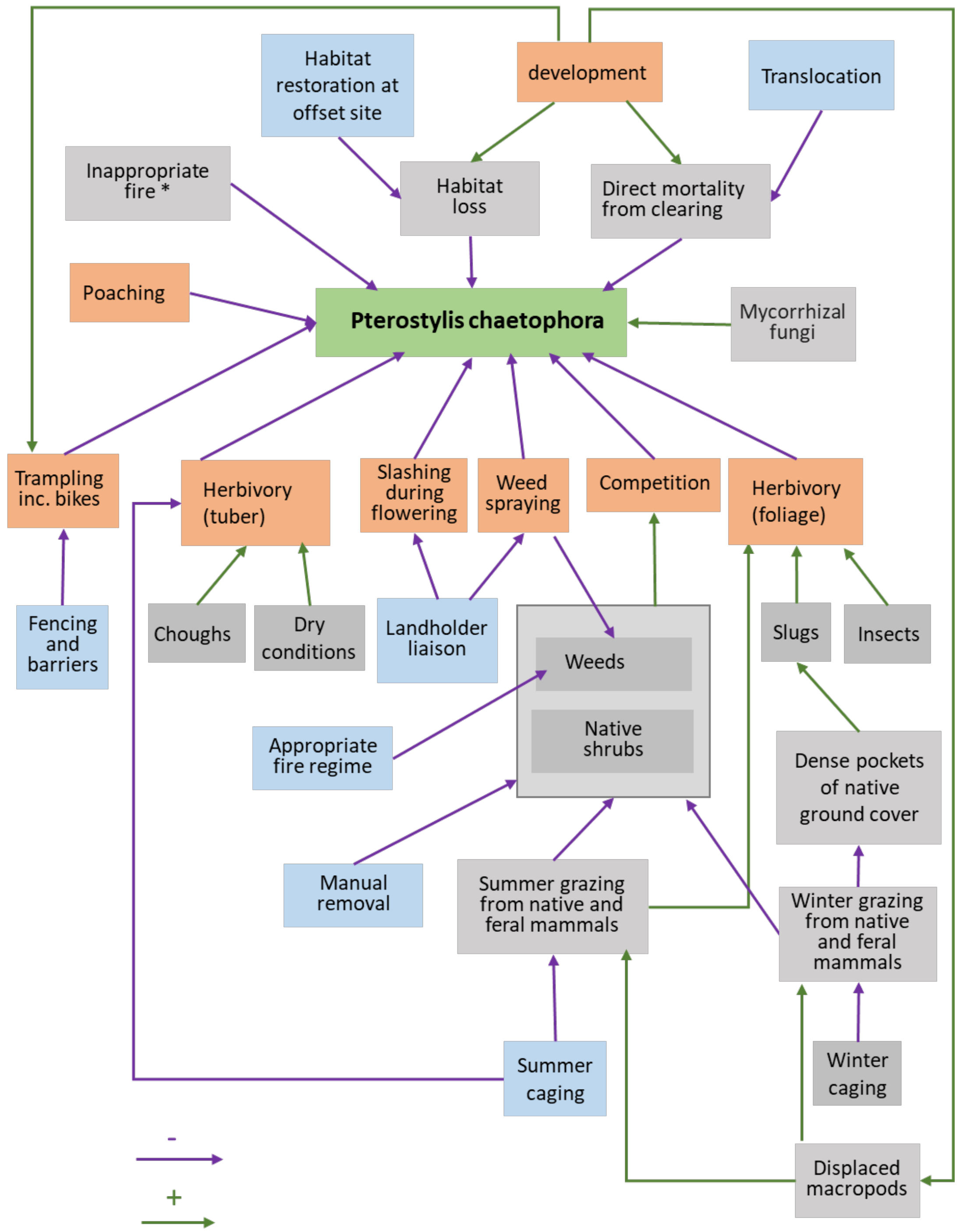


**Fig. 7:** Updated conceptual model for the tall rusthood

#### 4.3 *Tracking against modelled indicators*

The indicators selected in 2018 were number of individuals, proportion of chewed flowerheads, precent cover of weeds and disturbance. After five years, two major changes have been made to these. First the number of individuals is highly skewed, and variable between surveys depending on the how many individuals are visible. Based on this, the indicator for population will be changed to a grid-based occupancy model. Second, to reflect the difference in impact to species, weed cover has been split into ground cover weeds and shrubby weeds. Observed monitoring values for each indicator were provided by the species project coordinator (Fig 8). Additional information on each indicator has been extracted from the SoS monitoring database and is provided in Tables 11 - 14

The number of individual plants has not tracked as expected, with the values falling outside the estimated range of the both the managed and counterfactual unmanaged scenario. Percent weed cover for shrubby weeds responded as expected to management, however the coverage of groundcover weeds is higher than predicted if management actions had been fully implemented. Proportion of chewed flowerheads is above expected levels, and reflects resourcing issues, which impacted the ability to get cages onsite at the optimal time can therefore be inferred that they species itself has responded as expected and the decline reflects changes in threats (predations by choughs, sub-optimal caging due to lack of resources and COVID-19, and an increase in ground cover weeds.

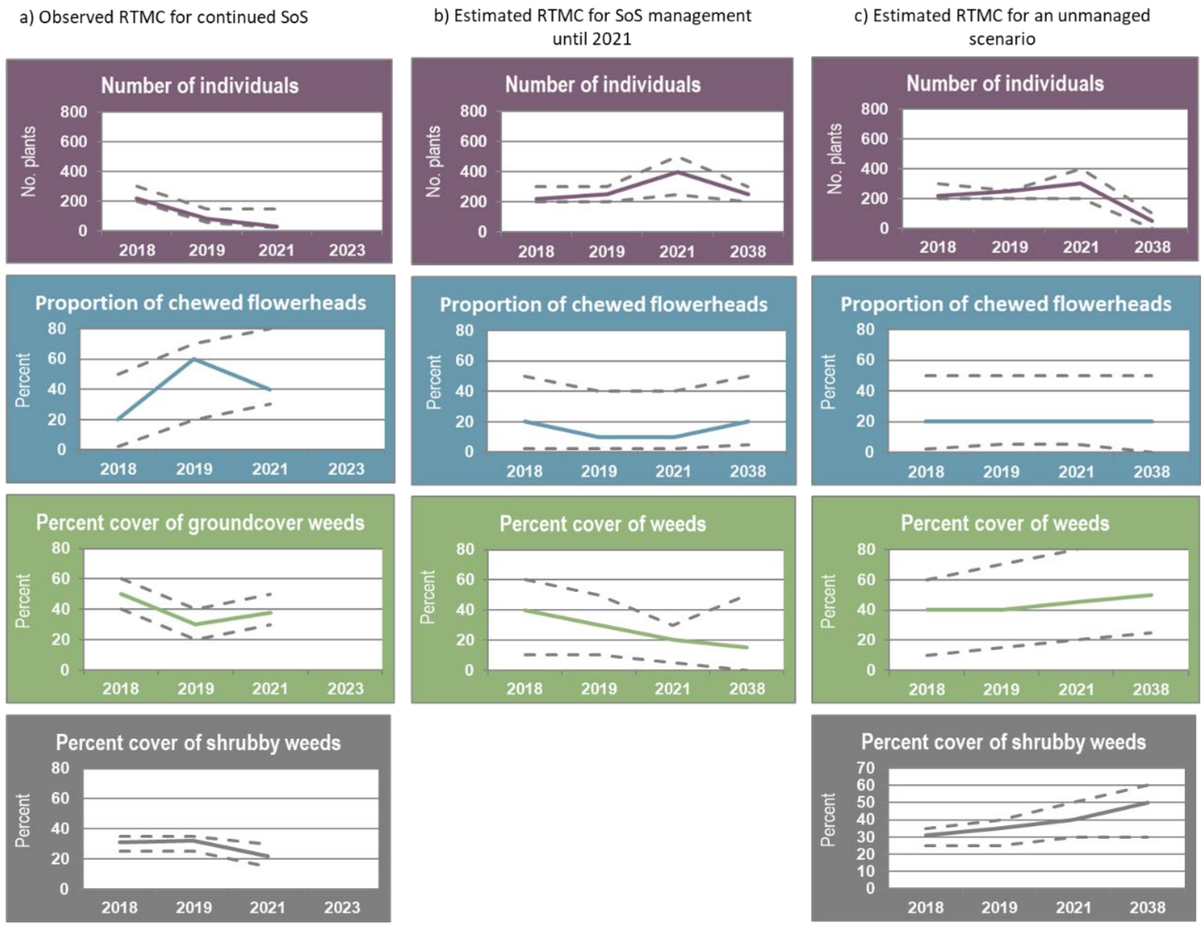


**Fig. 8:** Response to management curves for the 2018 tall rusthood for a) Observed RTMC b) Estimated RTMC for SoS management until 2021 c) Estimated RTMC for an unmanaged scenario. Values provided by Paul Hillier based on site monitoring data.

**Table 11:** Protection from grazing at the Black Creek Site for the tall rusthood orchid. Values extracted from SoS management database

| Financial Year | Monitoring Results | Confidence | Estimated Trend |
|---|---|---|---|
| 2018-2019 | Six cages were deployed and protected 45 individual plants. The grazing successfully protected plants from grazing with 87% reaching the flowering stage, compared to just 39% managing to flower within uncaged monitoring sites. Unfortunately only four plants progressed to seed production within two of the cages. Orchids within all cages began to develop capsules but towards the end of the monitoring period most withered in the dry conditions. | High | Not determined |
| 2019-2020 | A total of 81 plants were protected within eight cages. Due to the prevailing drought conditions, only four plants persisted through to seed production. The cages were also checked in August 2020 and a count of 84 emerging plants was made. This shows that whilst the dry conditions largely prevented seed production, the cages were successful in protecting the plants during a season where the threat from grazing and consumption by white-winged choughs was high. | Moderate | Stable |
| 2021-2022 | A total of 59 plants were protected by cages. However, non-emergence of orchids limited opportunities for protecting more flowers. There was no loss of from grazing upon caged plants. | High | Stable |
| 2022-2023 | Not applicable | Not applicable | Not determined |

**Table 12:** Weed coverage for the Black Creek Site of the tall rusthood orchid. Values extracted from SoS management database coverage

| Financial Year | Monitoring Results | Confidence | Estimated Trend |
|---|---|---|---|
| 2017-2018 | Three priority weed management areas have been identified, covering a total of 1.4 hectares. | High | Not determined |
| 2019-2020 | Between the monitoring events of 2018 and November 2019, the average exotic ground cover density had been reduced from 54% to 19%. The average exotic shrub layer density was largely unchanged at 31% in 2018 and 32% in 2019 and no exotic tree species have been recorded. Approximately 0.54 ha of additional hand weeding occurred in April 2020 around the known populations and significant't reduced the exotic ground and shrub layer. | Moderate | Not determined |
| 2020-2021 | The average exotic ground cover density within weed monitoring plots has reduced from 54% to 19% to 16% in 2018, 2019 and 2020 respectively. The average exotic shrub layer density has reduced from 32% in 2019 to 2% in 2020 and no exotic tree species have been recorded. Approximately 0.13 ha of additional hand weeding occurred in March 2021 around the known populations which further reduced the exotic ground and shrub layer. | Moderate | Decreasing |
| 2021-2022 | The average exotic ground cover density within weed monitoring plots has increased to 57%. The average exotic shrub cover density within weed monitoring plots has increased to 22%. Of the exotic species that are identified as a particular threat to Pterostylis chaetophora, the average ground and shrub covers are 17% and 21% respectively. | Moderate | Increasing |
| 2022-2023 | Not applicable | Not applicable | Not determined |
| 2018-2019 | All weeds surrounding known populations were removed. Weed monitoring plots were established prior to weed removal and these will be used to track the success of weed removal and track the severity of any weed reestablishment that may occur. | High | Not determined |

**Table 13:** Disturbance at the Black Creek site for the tall rusthood orchid. Values extracted from SoS management database

| Financial Year | Monitoring Results | Confidence | Estimated Trend |
|---|---|---|---|
| 2018-2019 | The use of a wildlife camera confirmed that motorbikes were still using the track. However, the addition of signs worked to reduce the threat and greater use of an alternate track was evident. Despite the ongoing use of motorbikes, monitoring of the track-side population confirmed that this population has not been impacted. A maximum of 20 plants were counted in the track-ide monitoring plot, compared to 13 counted in 2017. | High | Stable |
| 2019-2020 | No disturbance to plants or habitat was recorded within the site. | High | Stable |
| 2020-2021 | No disturbance to plants or habitat was recorded within the site. | High | Stable |
| 2021-2022 | No disturbance to plants or habitat was recorded within the site. | High | Stable |
| 2022-2023 | Not applicable | Not applicable | Not determined |

* in 2019 Disturbance to known populations has been managed through the construction of fencing and bollards at entry locations, and signage at one key location, which encourages motorbikes to use an alternative trail.

**Table 14:** Observed number of tall rusthood individuals at the Black Creek Site. Values extracted from SoS management database

| Financial Year | Monitoring Results | Confidence | Trend |
|---|---|---|---|
| 2019-2020 | A total of 30 individual plants were recorded within monitoring plots. | High | Decreasing |
| 2020-2021 | A total of 44 individual plants were recorded within monitoring plots. | Moderate | Not determined |
| 2021-2022 | A total of 11 individual plants were recorded within monitoring plots. The reduced count is believed to be a result of below average winter rainfall and not an actual decline in the population. The total population estimate is 120-200 individuals and has therefore not yet reached the long-term population target. | Moderate | Not determined |
| 2022-2023 | No plants were detected within monitoring plots. The reduction is believed to be due to the wet weather causing dormancy in this species. | Moderate | Not determined |

### 5 *Grevillea caleyi* (Caley's grevillea)

Contributors: Erica Mahon[1]

**Role/Affiliations:** [1] Species Project Coordinator, Ecosystems and Threatened Species – Greater Sydney Branch, Saving our Species Program, NSW Department of Environment, Energy, Climate Change and Water

#### 5.1 Summary

Caley's Grevillea (*Grevillea caleyi*) is a well understood, and well monitored species found within a limited 70-hectare range. In 2018, the critical management actions implemented for the project included: closure of tracks, weed control, fire management, education and potentially translocation. An offsite seed bank was also being created and maintained. The species was not affected by the black summer bushfire in 2019, although was affected by drought. While money was allocated to implement track closures, these were not approved by the area manager. Access was lost at one privately owned priority site, where weeds were self-managed. Phytophthora has now been detected at some sites, but it is not yet known whether Caley's Grevillea is susceptible to the pathogen. Since 2018, translocation has been successful, including of cuttings.

#### 5.2 Updates to the conceptual model

Several updates were made to the conceptual model (Fig 9).

- The definition of seed predation was expanded to include weevils, wallabies, and rats.
- Translocation is now considered a plausible, rather than potential management action against habitat loss (i.e. it is no longer represented with a dotted line). It is also linked to education, as one of the translocation sites is a school yard, creating an opportunity for student engagement. The link between translocation and improvement (or reduced decline) in genetic diversity has been added.
- Genetic diversity has been upgraded from a dashed to a solid line to reflect the fact we are now more certain that it's important.
- The definition of "Years since last fire" in model 2 has been refined to incorporate that this is years after some minimum threshold.
- The program is following research on odour-blocking deterrents to prevent over-grazing by native herbivores after fires. The feasibility of this is uncertain, but promising.
- Drought has been added as an explicated factor affecting post-fire rainfall.
- Phytophthora has now been detected at some sites, but it is not yet known whether Caley's Grevillea is susceptible to the pathogen. This has therefore been downgraded from a solid to a dotted line, as there are currently no plants showing signs.

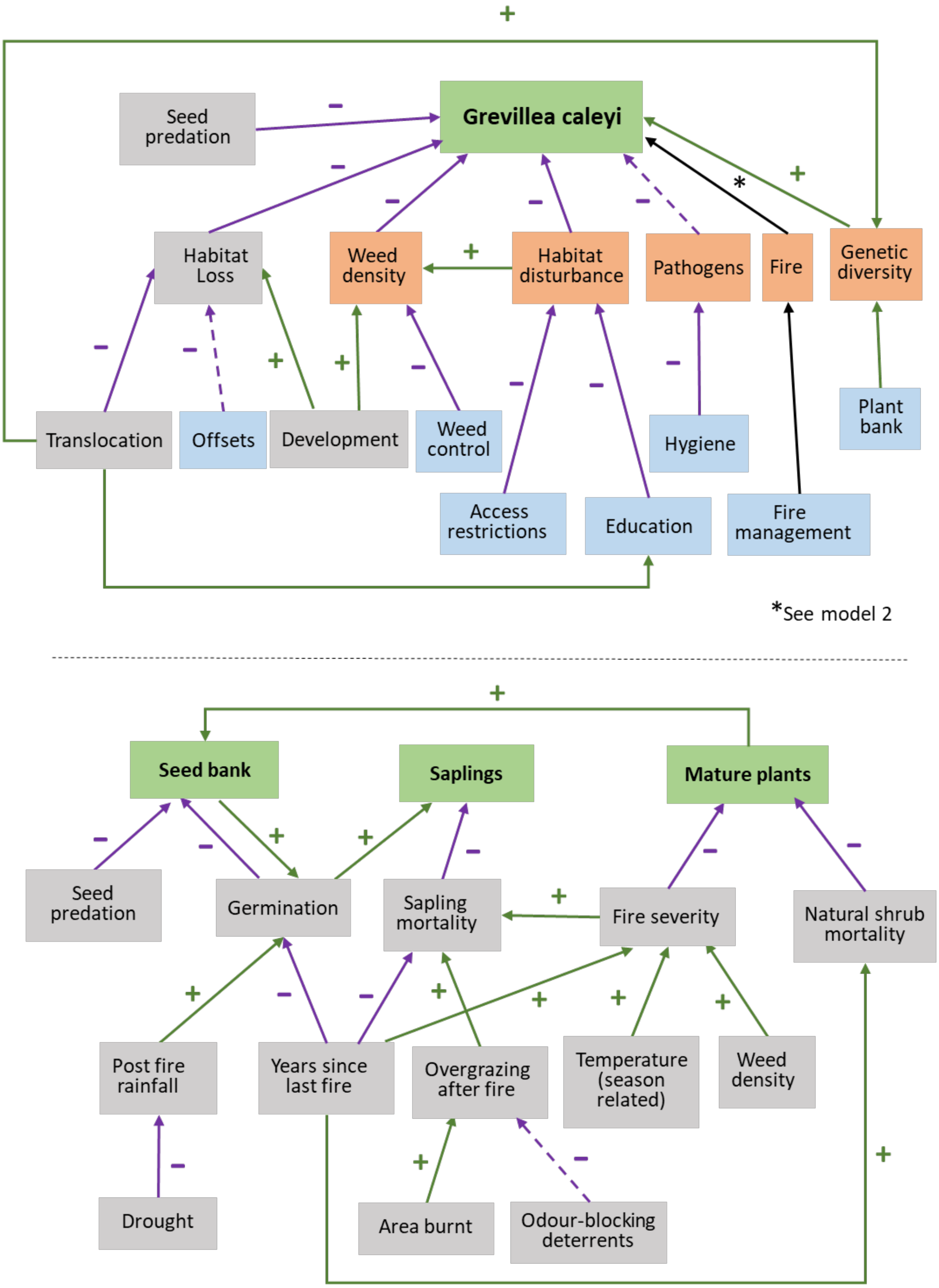


**Fig. 9:** Updated conceptual model for Caley's Grevillea

#### 5.3 *Tracking against modelled indicators*

The indicators selected in 2018 were post-fire seedling abundance, habitat area, weed density and relative disturbance from illegal tracks (Fig. 10).

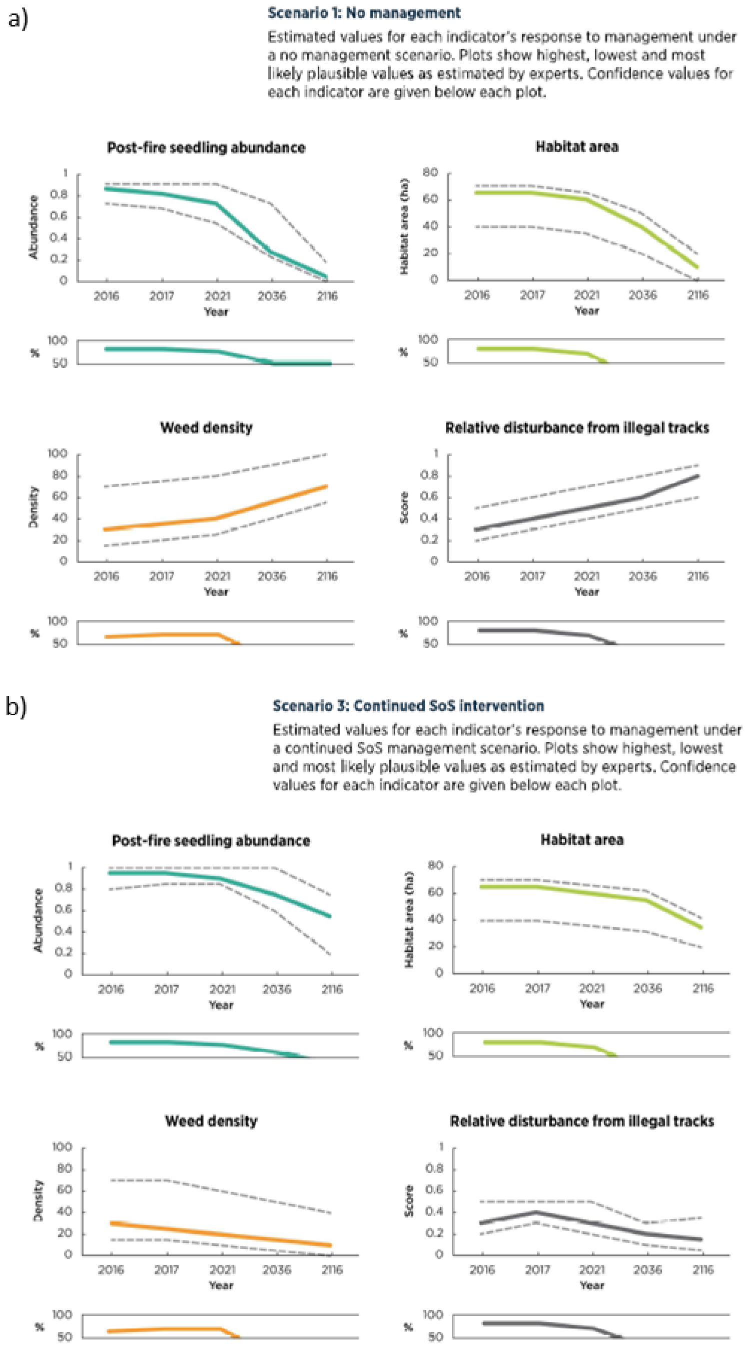


**Fig. 10:** Original indicators and estimated response to management curves for the 2018 Caley's Grevillea case study for an a) SoS managed scenario and b) a counterfactual unmanaged scenario

Post-fire seedling abundance was monitored, and a better-than-expected trend was observed (information provided by SoS species coordinator). Weed density sites that continued to be managed under SoS is within expected limits. At the site that is now self managed, the values for weed density are closer to those in Scenario 1 (no management). Habitat area remains stable (Fig. 11), although a road is being planned through one site, and the estimated loss is still expected to eventuate. Relative disturbance from illegal tracks has increased, as track closures were not approved (Scenario 1). This has likely had an impact on seedlings, despite the current higher than expected numbers observed.

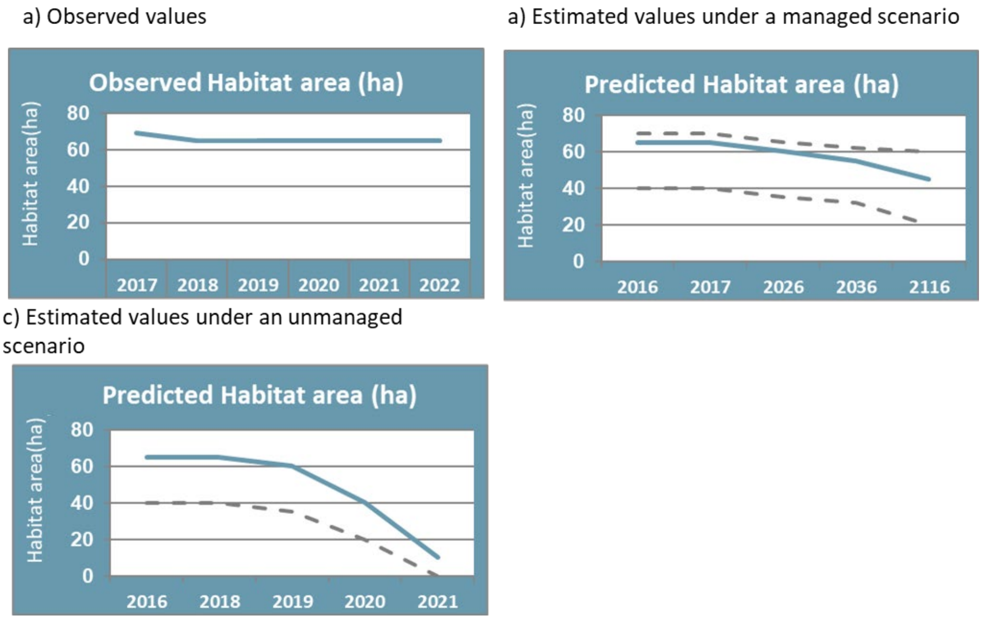


**Fig. 11:** Habitat amount for Caley's Grevillea case study for a a) observed values, b) estimated values for an SoS managed scenario and c) estimated values for a counterfactual unmanaged scenario

Additional qualitative information was also available from the SoS monitoring database (Tables 15-17)

**Table 15:** Observed habitat area for Caley's Grevillea. Data sourced from the SoS management database

| Year | Indicator | Description |
|---|---|---|
| 2017-2018 | 69 ha across 23 remnants. | |
| 2018-2019 | 64 ha. | Some small loss of habitat from recreational impacts but these may be able to be rehabilitated should the track be removed. |
| 2019-2020 | 64 ha | Habitat remaining is estimated as 64 ha, including buffer areas between patches. |
| 2020-2021 | 64.78 hectares. | The habitat area, including buffers, is 64.78 hectares. |
| 2021-2022 | 64.78 | Habitat area, including buffers, is 64.78 ha |
| 2022-2023 | 64.78 | No significant loss of habitat area, although minor areas were impacted through recreational activities. The habitat area is 64.78 hectares. |

**Table 16:** Fire management records for SoS sites managed for Caley's grevillea. Data sourced from the SoS management database

| Year | Description |
|---|---|
| 2017-2018 | Seedling recruitment (approximately 30) at Substation at Kamber Road following Hazard Reduction burn at the site. |
| 2018-2019 | Not applicable |
| 2019-2020 | No new fire event. Sites assesses have appropriate fire regime. Not all site assessed. |
| 2020-2021 | A number of burns led to good recruitment. No wildfires occurred so no areas have too frequent fire regimes. Some areas need fire as they have not been burnt or are long unburnt. |
| 2021-2022 | No wildfires occurred so no areas have too frequent fire regimes. Some areas need fire as they have not been burnt or are long unburnt. Planning with the University of Sydney for herbivory odour research. |
| 2022-2023 | No wildfires occurred so no areas have too frequent fire regimes. Some areas need fire as they have not been burnt or are long unburnt. No prescribed burns were recorded. |

**Table 17:** Disturbance records for SoS sites managed for Caley's grevillea. Data sourced from the SoS management database

| Year | Description |
|---|---|
| 2017-2018 | Disturbance of habitat is occurring from illegal track in Ku-ring-gai Chase and Garigal National Parks. Illegal track within the Ryland Remnant was mapped. |
| 2018-2019 | No new tracks were reported. The existing illegal track at the Ryland Track is still is use and there is evidence of small additional impacts from additional clearing e.g. to go around fallen tree, create a parallel path around a natural feature. |
| 2019-2020 | Disturbance from recreational and neighbours occurred at a number of remnants. |
| 2020-2021 | Disturbance to habitat occurred from recreational use of remnants such as mountain biking. |
| 2021-2022 | Recreational users continue to impact habitat areas. |
| 2022-2023 | The devices for this monitoring activity were purchased and delay delivered. Therefore, the actual monitoring activity hasn't been conducted. |

### 6 Hoplocephalus bungaroides (Broad-headed snake)

Revisited Case Study 6

Contributors: Meagan Hinds[1], Simon Lee[1], Caren Taylor[1], Ross Goldingay[2]

**Role/Affiliations:** [1] Species Project Coordinator Saving our Species, Ecosystems and Threatened Species – Greater Sydney Branch, Saving our Species Program, NSW Department of Environment, Energy, Climate Change and Water and [2] Species Expert, Southern Cross University

#### 6.1 Summary

Critical management actions were carried out at all three SoS sites, with additional context summarised in Table 18.

**Table 18:** Site summaries for the three broad-headed snake SoS management sites

| SoS site | Site summary |
|---|---|
| Morton National Park | Different habitat to the other two sites, comprising exposed cliffs rather than patchy rocky outcrops and surveys suggest the site likely has a higher density of snakes, compared to the other two sites. The group suggested that snakes may have been impacted by fires (from two years prior), with a downward trajectory observed. Additionally, threat monitoring data suggests an increase in disturbance which may also be contributing to the observed trajectory. |
| Royal National Parkland | Management was implemented as outlined in the species conservation strategy. The action to reduce the amount of human disturbance by closing trails appeared to result in the expected increase in population. Monitoring is only required every two years, as the population appears to be stable. |
| Woronora Plateau | Management has continued to reduce the disturbance, however the population is showing an unexplained downward trajectory. The group suggested reduced availability of crevices (exacerbated by high rainfall) could affect animals taking shelter under rocks, and the suitability of the site for the species |

#### 6.2 Updates to the conceptual model

The contribution of new experts, as well as updates in our knowledge, have led to numerous improvements to the original conceptual model. In particular, the scope of the model has been expanded to make it more applicable to sites outside of Morton National Park, e.g by including a link from trees to canopy cover. The following updates have been made to the conceptual model (Fig. 12):

- The model notation has been simplified to include juveniles as adults in the same block.
- Legal trade & Registry of licenced keepers has been combined
- Gecko poaching and snake poaching combined into reptile poaching. The link from reptile poaching to geckoes has been removed as it isn't considered a major factor in gecko abundance
- Suitable rocks changed to exposed rocks, to reflect the relationship with vegetation.
- Crevices has been added, which are used by snakes for sheltering in and ambush hunting
- Feedback loop between reptile poaching and human disturbance has been added to show that increased human disturbance can increase poaching.

- Surveillance added, but needs to be linked in
- Goats have been removed. Although they can crack rocks, the impact is not considered substantial or a critical threat.
- The link from mature trees to canopy cover has been included. It has been represented as a dotted line to indicate some uncertainty as to whether the use of trees and exposed rocks overlaps geographically. i.e. the mature trees used my snakes for tree hollows may not be in the same area as the exposed rocks.
- Small-eyed snakes were removed as they have been found to co-exist in the same area
- Rain reduces the availability of exposed rocks as leaf litter and debris will get stuck under the rocks, making them unsuitable.

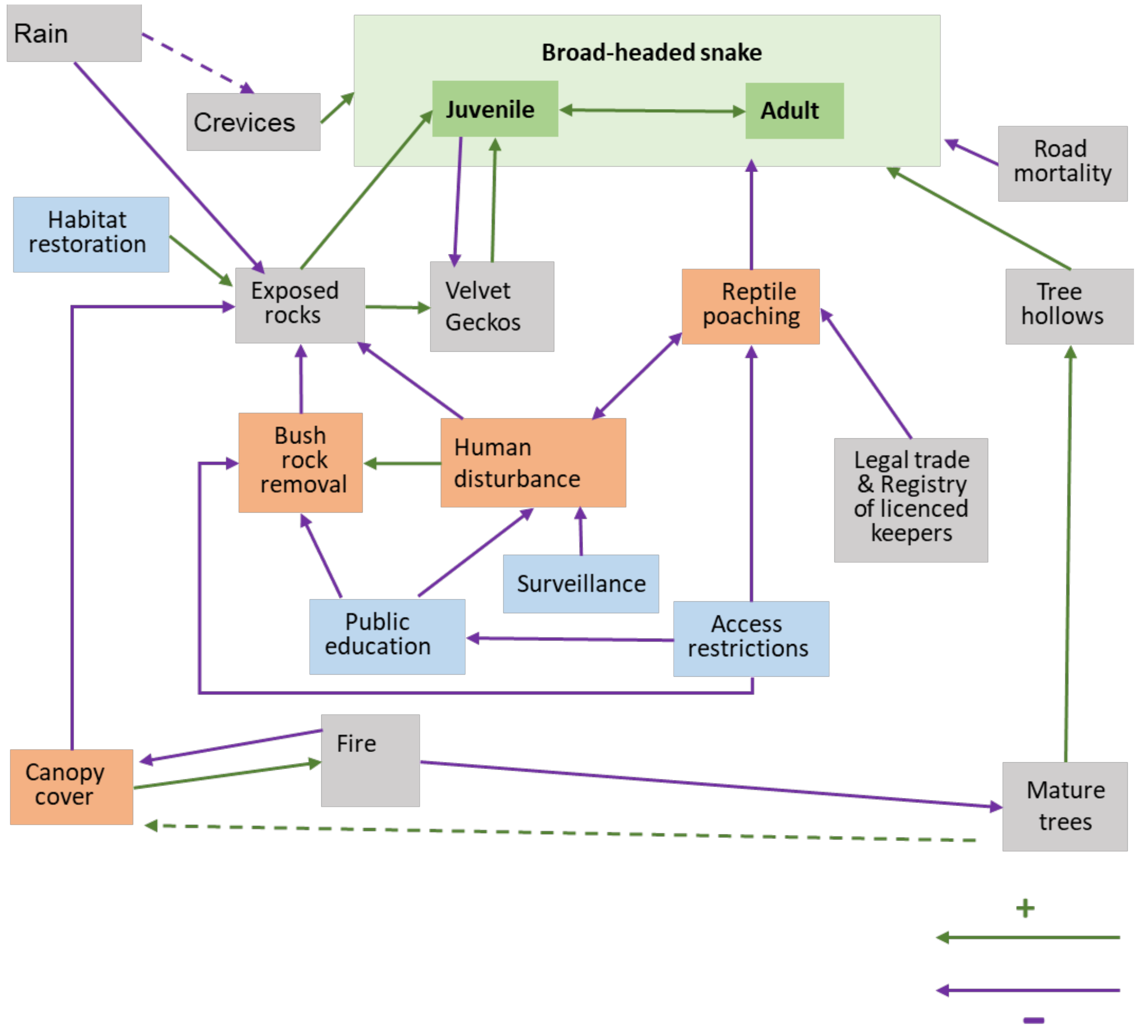


**Fig. 12:** Updated conceptual model for Broad-headed snake

#### 6.3 *Tracking against modelled indicators*

The broad-headed snake has three sites SoS sites. .The original estimates from 2018 (Fig. 13) are most closely associated with Morton National Park. There are some key differences between sites which affect the way in which the population can be measured. In Woronora Plateau and Royal National Parkland, the trees used by the snakes are closer to the basking platforms than at the Morton National Park Sites. .This means adult snakes are likely to utilise the trees more than at Morton. The snakes found under rocks at the first two sites are therefore primarily juveniles, compared to adult snakes at Morton. The result of this difference is that while capture-recapture monitoring is appropriate for Morton, occupancy is more suitable for Royal National Park and Woronora Plateau, where adults are less likely to be found.

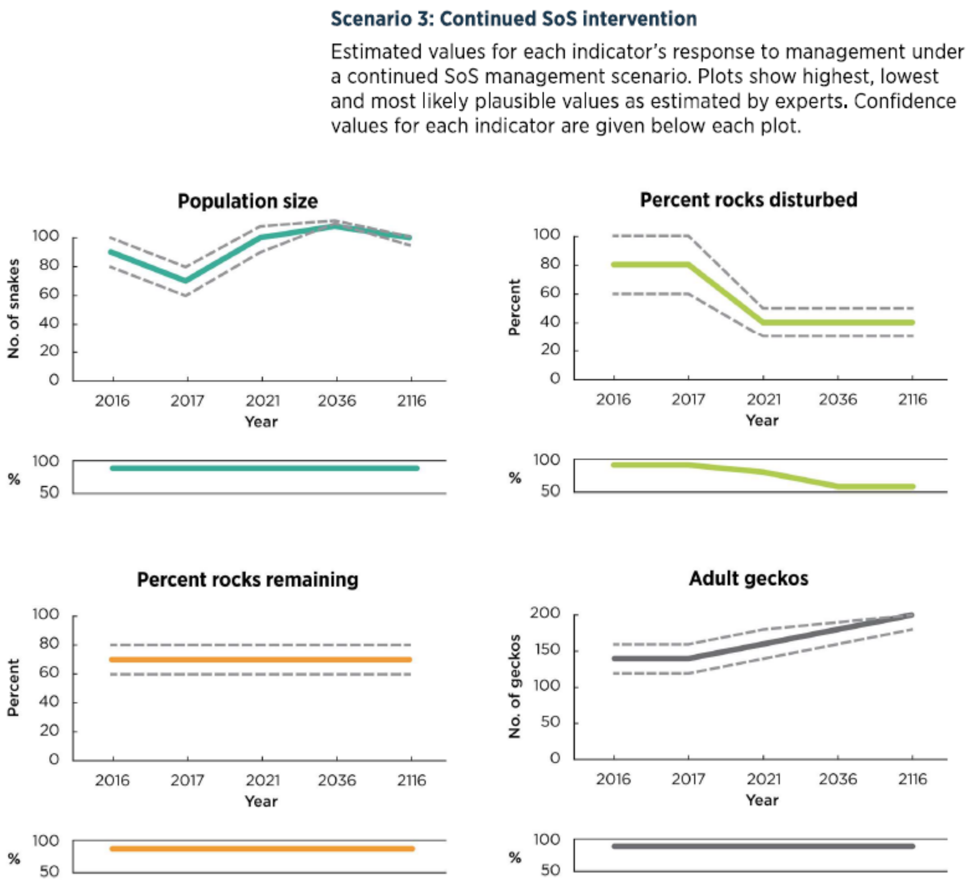


**Fig. 13:** Original indicators and estimated response to management curves for the 2018 Broad-headed snake case study.

Three of the indicators were monitored at most sites; Occupancy, extent of bush rock disturbance, and likelihood of illegal collection (Table 19 – Table 21). Adult geckos are no longer considered as a suitable indicator, as a large gecko population is often associated with an absence of broad-headed snakes. They are however still worth monitoring as an indicator of habitat quality. The size of the population is not a feasible indicator outside of the Morton site, as capture-recapture monitoring is not feasible at Royal National Park or Woronora Plateau.

Based on discussions with site experts, occupancy at Royal National Park is presumed to have remained stable at around 50%, allowing for the uncertainty surrounding monitoring data. Woronora Plateau maintains a stable occupancy level of 30% for the last few years, however this is down from 70% in 2017. The population at Morton has fluctuated periodically in response to poaching events. Observed values are given in Fig 14.

a) Observed Occupancy

a) Estimated population size for a managed scenario

a) Estimated population size for a counterfactual unmanaged scenario

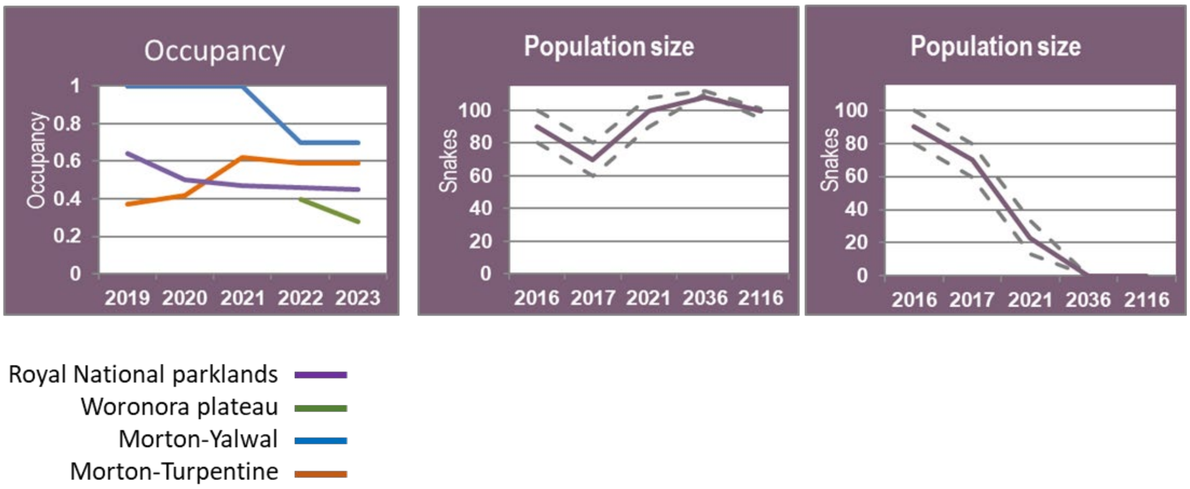


**Fig. 14:** Values for a) observed occupancy, b) predicted population size for an SoS managed scenario and c) population size for a counterfactual unmanaged scenario

**Table 19** Occupancy of broad-headed snake across the monitored SoS sites

| Year | Royal National Parkland | Woronora Plateau | Morton-Yalwal | Morton-Turpentine |
|---|---|---|---|---|
| 2018-2019 | 0.64 ± 0.20 | - | 1 | 0.37 +-0.006 |
| 2019-2020 | - | - | 1 | 0.42 +-0.17 |
| 2020-2021 | 0.47 +-0.1 | - | 1 | 0.62 +-0.17 |
| 2021-2022 | - | 0.4 +- 0.1 | 0.7 +- 0.12 | 0.59 +-0.15 |
| 2022-2023 | 0.45±0.06 | 0.28 ± 0.09 | 0.7 +- 0.12 | 0.59 +-0.15 |

**Table 20** Extent of bush rock habitat disturbed for broad-headed snake across the monitored SoS sites

| Year | Royal National Parkland | Woronora Plateau | Morton-Yalwal | Morton-Turpentine |
|---|---|---|---|---|
| 2017-2018 | 7 of 26 | 1 of 42 | - | - |
| 2018-2019 | 2 of 27 | 7 of 31 | - | - |
| 2019-2020 | - | 8 of 31 | - | - |
| 2020-2021 | 0 of 26 | 2 of 31 | - | - |
| 2021-2022 | - | 6 of 31 | - | - |
| 2022-2023 | - | - | - | - |

**Table 21** Likelihood of illegal collection of broad-headed snakes across the monitored SoS sites, inferred from bush rock habitat

| Year | Royal National Parkland | Woronora Plateau | Morton-Yalwal | Morton-Turpentine |
|---|---|---|---|---|
| 2017-2018 | - | - | - | - |
| 2018-2019 | - | - | - | - |
| 2019-2020 | - | - | Low | Moderate |
| 2020-2021 | - | - | Low | Moderate |
| 2021-2022 | - | - | Moderate | Moderate |
| 2022-2023 | - | - | Moderate | Moderate |
