## Supplementary material for "Evaluating a structured expert elicitation approach for adaptive conservation: Lessons from five years in practice": S2 Table - Conceptual diagram updates

**Supplementary S2 Table.** Key changes to conceptual diagrams for each of the six review case studies

| Case study | Key changes to model |
| --- | --- |
| Long-nosed potoroo | ● The link between deer/pigs and long-nosed potoroo populations now considered unlikely<br>● Habitat quality and structure added as a limiting factor to predation<br>● The importance of cats, wild dogs, foxes and pigs differs at each site. |
| Large bent-winged bat | ● Windfarms now a confirmed rather than uncertain threat<br>● Distance of turbines to roost has been included<br>● Presence of cats is no longer consider a key threat |
| Mahoney's toadlet | ● Connectivity between core and peripheral sites is now represented<br>● Development and sand mining are not relevant for the population in a State protected area<br>● Habitat suitability is driven primarily by geology and soil<br>● Chytrid is now considered a potential threat (previously uncertain), as it has been identified in some sub-populations.<br>● Water extraction is a possible threat at some sites<br>● Gambusia are now considered a potential threat (previously uncertain) |
| Tall rusthood | ● Grazing has been separated into growth-period (Aug-Jan) and dormant period (Feb-July) grazing<br>● The new threat of nearby development has been added<br>● Herbivory from choughs identified as new threat |
| Caley's grevillea | ● Translocation is now considered a plausible, rather than potential management action<br>● Genetic diversity now a certain factor<br>● odour-blocking deterrents to prevent over-grazing by native herbivores after fires a potential management action.<br>● Drought has been added as an explicit factor affecting post-fire rainfall<br>● Phytophthora has now been detected at some sites, but it is not yet known whether Caley's grevillea is susceptible to the pathogen |
| Broad-headed snake | ● Legal trade & Registry of licenced keepers has been combined<br>● Goats have been removed<br>● The link from mature trees to canopy cover has been included<br>● Small-eyed snakes were removed as they have been found to co-exist in the same area<br>● Rain reduces the availability of exposed rocks as leaf litter and debris will get stuck under the rocks, making them unsuitable. |
