## Supplementary material for "Evaluating a structured expert elicitation approach for adaptive conservation: Lessons from five years in practice": S3 Table - Observed v predicted vaules

**Supplementary S3 Table:** Observed and predicted (with and without management ) values for case study species from 2016 to 2024 in New South Wales. Decimal points reflect the available precision of the data

| | Year | | | | | | | | |
|---|---|---|---|---|---|---|---|---|---|
| | **2016** | **2017** | **2018** | **2019** | **2020** | **2021** | **2022** | **2023** | **2024** |
| | (min-max, confidence) | (min-max, confidence) | (min-max, confidence) | (min-max, confidence) | | (min-max, confidence) | | | |
| **Long-nosed potoroo (Proportion of camera traps with potoroos)** | | | | | | | | | |
| Observed | | | 0.50 | | 0.50 | | 0.55 | | 0.55 |
| Predicted with management | 0.10 (0.05-0.15, 80%) | 0.1 0 (0.05-0.16, 80%) | | 0.09 (0.04-015, %) | | | | | |
| Predicted without management | | 0.1 0 (0.04-014, 80%) | | 0.08 (0.02-0.12, %) | | | | | |
| **Large bent-winged bat (1000's of bats at roost site)** | | | | | | | | | |
| Observed | | 23.7 | 18.0 | 18.4 | 16.9 | 19.2 | 23.0 | | |
| Predicted with management | 17.8 (17.6-18.0, 95%) | | | | | 21.5 (21.0 - 22.0, 95%) | | | |
| Predicted without management | | | | | | 17 (10.0 - 21.0, 80%) | | | |
| **Tall rusthood (Numer of orchids - modelled estimates)** | | | | | | | | | |
| Observed | | | 220 (200-300, 60%) | 80 (57-150, 95%) | | 30 (23-150, 95%) | | | |
| Predicted with management | | | | 250 (200-300, 50%) | | 400 (250-500, 50%) | | | |
| Predicted without management | | | | 250 (200-300, 50%) | | 300 (200-400, 50%) | | | |
| **Caley's grevillea (Hectares of habitat)** | | | | | | | | | |
| Observed | | 69 | 65 | 65 | 65 | 65 | 65 | | |
| Predicted with management | 65 (70-40, 80%) | 65 (70-40, 80%) | | | | 60 (35-65, 80%) | | | |
| Predicted without management | | 65 (70-40, 80%) | | | | 60 (35-60, 75%) | | | |

| | Year | | | | | | | | |
|---|---|---|---|---|---|---|---|---|---|
| | **2016** | **2017** | **2018** | **2019** | **2020** | **2021** | **2022** | **2023** | **2024** |
| | (min-max, confidence) | (min-max, confidence) | (min-max, confidence) | | | (min-max, confidence) | | | |
| **Broad-headed snake (Number of snakes)** | | | | | | | | | |
| Observed (Royal) | | | | 0.64 | 0.50 | 0.47 | 0.46 | 0.45 | |
| Observed (Woronora) | | | | | | | 0.40 | 0.28 | |
| Observed (Morton-Yalwal) | | | | 1.00 | 1.00 | 1.00 | 0.70 | 0.70 | |
| Observed (Morton-Turpentine) | | | | 0.37 | 0.42 | 0.62 | 0.59 | 0.59 | |
| **Broad-headed snake (Occupancy, percent of site occupied - All sites)** | | | | | | | | | |
| Predicted with management | 90 (70-80, 90%) | 70 (60-80, 90%) | | | | 100 (90-108, 90%) | | | |
| Predicted without management | | 70 (60-80, 90%) | | | | 23 (13-33, 90%) | | | |
