## Supplementary material for "Evaluating a structured expert elicitation approach for adaptive conservation: Lessons from five years in practice": S4 - Multi-species case studies

#### Supplementary S4: Additional case studies

Helen J Mayfield[1,2], James Brazill-Boast[3], Mick Andren[3], Michiala Bowen[3], Adam Fawcett[3], Trent Forge[3], Luke Foster[3], Ross Goldingay[4], Paul Hillier[3], Meagan Hinds[3], Simon Lee[3], Erica Mahon[3], Martine Maron[1,2], Doug Mills[3], Thomas Rowell[3], Stephanie Stuart[3], Caren Taylor[3], Grant Webster[5,6], Nicole Hansen[3]

[1] School of the Environment, The University of Queensland, Brisbane, QLD Australia
[2] Centre for Biodiversity and Conservation Science, The University of Queensland, Brisbane, QLD Australia
[3] Saving our Species Program, New South Wales Department of Climate Change, Environment and Water, New South Wales, Australia
[4] Southern Cross University, Lismore, New South Wales, Australia
[5] University of New England, New South Wales, Australia
[6] Institute for Applied Ecology, University of Canberra, Bruce, Australian Capital Territory, 2617, Australia

### 1 Mount Kaputar Land Snail & Slug Threatened Ecological Community

**Adam Fawcett**

#### 1.1 Introduction

Mount Kaputar Land Snail & Slug TEC occurs within the Nandewar Ranges near Narrabri, centred on Mount Kaputar NP. The TEC was the first of its kind listed (possibly worldwide) due to the high diversity of snail species within an island refuge. The TEC consists of at least 12 native land snail species and one giant pink slug occurring within the high-altitude areas (>1000m above sea level) of Mount Kaputar National Park of the Nandewar Ranges. Below this altitude, the dry rainforest communities act as important refuge areas down to ~500m above sea level. As at 2022, Monitoring data of the component species within the TEC is limited, have only being undertaken in the past 12 months. Future monitoring will help inform and improve our understanding of the system.

A large proportion of the TEC is reserved within Mount Kaputar National Park (MKNP). Access to the reserve is via a single main road leading to the summit. There are several additional fire trails within the reserve providing access to some areas but the majority of the reserve is declared wilderness and only accessible on foot. Facilities across MKNP include campgrounds, walking trails and lookouts. In addition, the summit of Mount Dowe includes a major telecommunications tower and associated infrastructure.

#### 1.2 Scope

The Mount Kaputar Land Snail & Slug TEC occurs over an area of ~163,000 ha within the Nandewar Ranges. The TEC exists at two sites, the Nandewar Ranges and East Horton. This site covers a range of landscape but the focus areas for the TEC includes the high elevation areas and the dry rainforest, as described above. The model refers to the entire distribution as the threats to the TEC are similar across both sites. The only additional factor influencing the East Horton site is habitat loss from land clearing and grazing.

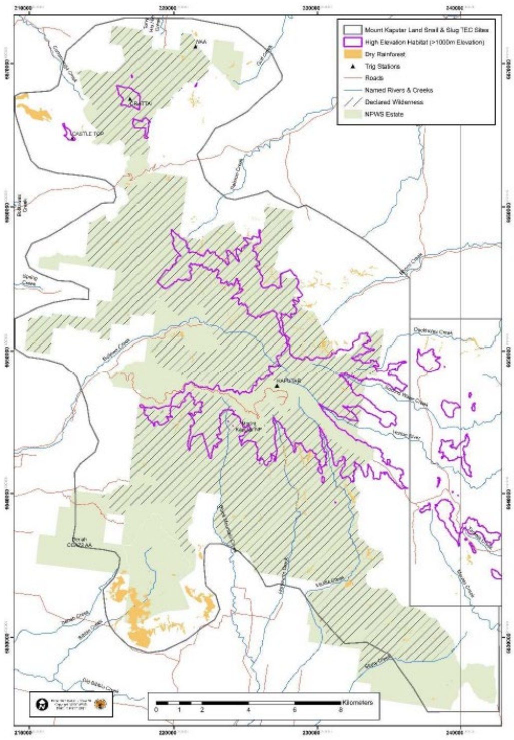


**Fig. 1:** Scope of the Mt Kaputar TEC

Current interventions in place include

- Pig control via trapping programs within sub-alpine areas and aerial shoot programs
- Goat control via aerial shoot programs, mustering and ground shooting
- Advice on fire management to limit the impacts of frequent fire, fire severity and ensure exclusion from dry rainforest communities. This targets fire planning and patch size for hazard reduction burns
- Advice on the construction of additional roads and paths as part of fire management planning but also infrastructure planning and maintenance.

#### 1.3 Narrative

- **Snail & Slug Species Richness and Diversity:** Surveys for the component species that make up the TEC use species richness from repeat site searches to understand changes in the community over time. Number of species detected feeds directly into species diversity and influences TEC quality. Species diversity (presence or absence of the component species) provide a measure of the complexity and quality of the overall TEC quality.
- **Habitat Quality:** Habitat quality, is influenced **by vegetation structure, leaf litter, ground moisture, rocks and logs/timber** Reduction in habitat quality negatively impacts TEC species richness and diversity.
- **Ground moisture** improves microhabitat conditions that influence habitat quality.as is influenced by climatic conditions.  It is improved with, leaf litter, logs and timber on the ground and is negatively impacted by drought.

- **Leaf litter** deteriorates with a deterioration in **vegetation structure** or removal / lack of **rocks**. Fire reduces leaf litter.
- **Logs/timber on ground:** Logs/timber on ground provides refuge habitat and increases habitat quality. While fire decreases available logs/timber on the ground, large logs can also provide refuge for snail species from low severity fire, reducing mortality from fire.
- **Rock:** Rock, both as outcrops and loose rock on the ground, provides refuge areas for snails and slugs, increasing habitat quality. It also provides refuge from trampling from feral animals, park users and livestock along roadsides & access trails and, as a refuge, also reduces mortality from fire.
- **Fire:** Fire has a negative effect on vegetation structure and directly impacts on individual snails and slugs through mortality from fire. The impact of fire is variable depending on fire frequency and severity. Increased fire frequency and severity is one of the symptoms of Climate Change.
- **Mortality from Fire:** Fire directly impacts on individual snails and slugs, reducing the species richness and diversity of the TEC. Fire related mortality can be decreased by the presence of rock and logs/timber on the ground.
- **Fire Management:** Fire management is aimed at reducing the severity and extent of fires but can result in the increased number of roads and paths within the reserve.
- **Roads and paths:** Construction of roads and paths impact on vegetation structure and increase the effects of mortality from trampling. As construction is often associated with fire management, fire has an increasing effect on this threat. Construction is also highly driven by increasing visitor numbers within the reserve and the need to improve infrastructure and assets accordingly.
- **Mortality from Trampling:** Mortality from trampling by feral animals, park users and livestock has a negative effect on the species richness and diversity of the TEC through direct loss of individuals and is exacerbated by the presence of roads and trails.
- **Drought:** Drought has a negative impact on vegetation structure and ground moisture while increasing the potential for fire frequency & severity. Drought is one of the symptoms of Climate Change.
- **Pigs:** Pigs negatively impact vegetation structure thru soil & vegetation disturbance and predate directly on snails & slugs. Pigs can be managed through **feral animal control.**
- **Goats:** Goats negatively impact on vegetation structure via grazing and trampling. Goats can be managed through **feral animal control.**
- **Black Rats:** Black rats negatively impact via predation on snails & slugs. This is not a considered a critical threat at this stage and more work needs to be completed to identify the importance of this potential threat.
- **Snail predation:** Snail predation by black rats and pigs decreases on species richness and diversity of the TEC.
- Several threats are specific to the Eastern Horton site. These are **land clearing** for grazing lands, **introduced weeds** and **grazing** by livestock. All of these affect habitat quality. Land clearing increases grazing and introduced weeds, and grazing also increases the weeds and trampling by livestock.
- **Vegetation structure:** The TEC is influenced by vegetation structure. Several snail species and the pink slugs are only found within the high-altitude communities (1000 metres above sea level), which includes the sub-alpine forest on the higher peaks and the open forests on the surrounding slopes. At lower altitudes, the dry rainforest communities provide the key habitat for the snail species. As the structure of these vegetation communities are influenced by several additional factors, a secondary model identifies TEC drivers of vegetation structure.

- **Drought:** Drought can cause direct mortality of vegetation and/or loss of canopy cover, affecting all three communities (dry rainforest, open forest and sub-alpine). Drought can also result in increased fire severity and frequency.
- **High fire severity: High f**ire severity negatively impacts on the structural components of both sub-alpine and open forest communities. High fire severity reduces the quality of both grass dominated and shrub dominated communities. High fire severity also results in a loss of canopy cover of dry rainforest communities which can potentially result in increased fire frequency.
- **Fire frequency:** Increased fire frequency increases fire severity while also increasing structural changes within vegetation communities. Increased fire frequency can also reduce the presence of a shrub layer within open forest communities, making them grassier over time.
- **Hazard Reduction Burns:** Hazard reduction burns are aimed at reducing the overall fuel hazard and protecting human life and property from the effects of unplanned wildfire. If not undertaken correctly (**Improperly executed**) there is a potential for a direct impact on dry rainforest communities and canopy cover. If conducted correctly (**Executed as planned**), hazard reduction burns may have a positive influence on vegetation mortality, by reducing the severity of fire over time.
- **Loss of Canopy Cover:** Loss of canopy cover occurs as result of mortality from fire and drought effects within dry rainforest communities. There is potential that it can also in turn increase fire frequency .
- **Structural complexity of High-altitude vegetation communities:** For the vegetation of the TEC above 1000m AHD, the communities tend to either be **grass dominated** (sub-alpine communities on high peaks) or **shrub dominated** (open forests on surrounding slopes). The structure of these communities is influenced by changes incurred by drivers of the TEC. Increased fire frequency will push shrub dominated communities towards grass dominated over time. High fire frequency can result in an increased shrub layer within normally grass dominated communities. Succession over time within these communities will result in
- the communities moving back toward their preferred state (either shrub or grass dominated), depending on further impacts from TEC drivers. Management of these communities aims to achieve a balance through implementing appropriate fire regimes for the respect vegetation community.

#### 1.4 Conceptual model

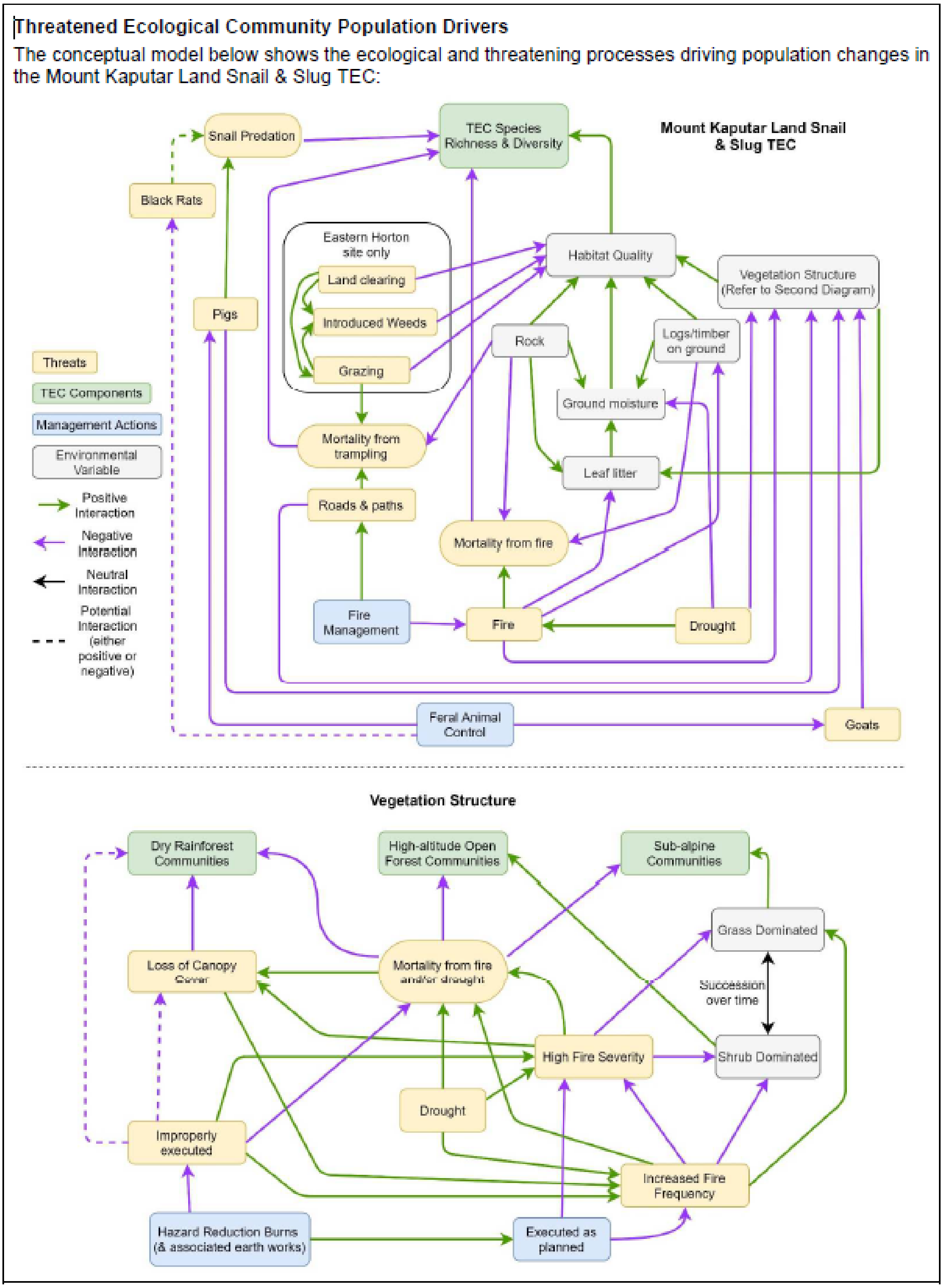


**Fig. 2:** Conceptual model for the Mount Kaputar Land Snail & Slug Threatened Ecological Community

#### Conceptual model definitions

| Snail & Slug Species Richness and Diversity | The number of different component species present at a site |
|---|---|
| Black Rats | Abundance of rats across both sites |
| Drought | Severity and length of droughts in a given time period |
| Dry rainforest communities | Condition and extent of the dry rainforest communities which provide the key habitat for the snail species at lower altitudes |
| Executed as planned | The percent of hazard reduction burns that are executed appropriately and as planned |
| Feral animal management | Intensity and scope of feral animal management (goats and pigs) |
| Fire management | Location and frequency of hazard reduction burns |
| Goats | Abundance of goats across both sites |
| Grass Dominated | Amount of area that is grass dominated |
| Grazing | Grazing pressure from livestock or feral herbivores |
| Ground moisture | |
| Habitat Quality | Suitability of the habitat for component species. All species have similar requirements |
| Hazard Reduction Burns | Frequency of managed, appropriately timed, low-intensity hazed-reduction burns. |
| High-altitude open forest communities | Condition and extent of High-altitude open forest communities which occur above 1000m |
| Introduced weeds | Amount of habitat with more than x% of introduced weeds |
| Improperly executed | The percent of hazard reduction burns that are executed inappropriately |
| Land clearing | Amount of habitat lost to clearing |
| Leaf litter | Depth? % coverage? |
| Logs/timber on ground | Distribution density of fallen logs for snails and slugs to hide under |

| | |
|---|---|
| Loss of Canopy Cover | Percentage of canopy cover |
| Fire | Intensity and frequency of fire events |
| High Fire severity | |
| Increased fire frequency | Number of fires in a given time period |
| Mortality from Fire | Number of individuals directly killed from a fire event (not directly measured) |
| Mortality from Trampling | Number of individual snails/slugs trampled by people |
| Roads and paths | |
| Rock | Distribution density of large, undisturbed rocks for snails and slugs to hide under |
| Shrub Dominated | Amount of area that is shrub dominated |
| Snail predation | Number of snails eaten |
| Sub-alpine communities | Condition and extent of sub-alpine forest which occurs on the higher peaks |
| Vegetation structure | Suitability of the vegetation structure as habitat for snail and slug component species |

### 2 Floyd’s Grass and Black Grass-dart Butterfly

Mick Andren

#### 2.1 Introduction

**Floyd’s Grass** *Alexfloydia repens* occurs in two discrete regions on the NSW north coast: Sawtell and Warrell Ck (Andren and Cameron 2012). SoS sites have been established at Pine Ck (within the Sawtell population and containing by far the largest amount of grass), Warrell Ck and Digger’s Headland (a small outlying ‘insurance’ population). **Floyd’s Grass is the key (obligate) habitat of the Endangered Black Grass-dart Butterfly** which has a separate SoS Monitoring Plan.

For Floyd’s Grass, The objective of the project in the short to medium term will be maintenance or increase in the extent of the grass. In the longer term, sea-level rise will begin to reduce the extent. At that point other options will need to be explored, such as translocation or propagating and selling Floyd’s Grass as an ornamental plant. Floyd’s Grass has already been translocated with success, so there are no urgent reasons to act on the sea-level rise threat at present (this is unlikely to change for several decades).

**The Black Grass-dart Butterfly** *Ocybadistes knightorum* occurs in two discrete regions on the NSW north coast: Sawtell and Warrell Ck (Andren and Cameron 2012). SoS sites have been established at Pine Ck (within the Sawtell population and containing by far the largest number of butterflies), Warrell Ck and Digger’s Headland (a small outlying ‘insurance’ population). The Floyd’s Grass SoS weeding project is also the main management action for the conservation of the butterfly.

#### 2.2 Scope

Floyd’s Grass occurs in discrete patches in a restricted geographical area and can therefore be accurately mapped (Andren and Cameron 2012). The first mapping exercise was in 2010, which mapped 32.6ha in 293 patches of varying size (Andren and Cameron 2012). From experience with this species, re-mapping once a decade is considered a suitable timeframe to detect significant changes in extent.  Three sites are included in SoS.

- The Pine Creek site is located south of Sawtell on the NSW north coast. The site includes low-lying swamp forest adjacent to the creek along a 7km section, beginning 3.5km from the creek mouth at Sawtell. This site contains 72% of all known Floyd’s Grass as it encompasses most of the largest patches.
- The Warrell Creek site is located between Nambucca Heads and Scotts Head on the NSW north coast. The site includes low-lying swamp forest adjacent to the creek along a 4km section, beginning about 7km south of the creek mouth at Nambucca Heads. It is likely that this population is disjunct from the main population at Sawtell.
- Digger’s Headland is located in Coffs Harbour and contains an atypical occurrence of Floyd’s Grass on the southern slope of the headland. It is well above sea level and therefore not at risk from sea-level rise.

-

#### 2.3 Narrative

- Floyds grass provide shelter for the butterfly at all life stages, and provides food for larvae
- Weed infestation. Habitat quality and extent can be greatly reduced by weed invasion, which can be managed by weed control
- Encroachment by native plants. This can occur to due canopy shading, increased exposure, vegetation succession, aggressive native invaders, post-fire regeneration, etc. Thinning of native plants is an option, but is not currently being considered.
- Fire. The best quality Floyd's Grass habitat occurs on peat soils. During very dry conditions, peat will burn continuously once a fire is started. This is even more critical if the fire occurs when there are no adult stages (butterflies) that can escape the fire. Protection from these intense peat fires is important for both species.
- Climate change will result in habitat loss. Floyd's Grass occurs directly above the king tide zone and most of it will be lost due to predicted sea level rise (Andren and Cameron 2014).
- Climate change will also increase the frequency of droughts, which negatively impacts Floyd's grass. Droughts and higher temperatures will also increase desiccation in butterfly larvae and eggs, leading to increased mortality, as well as reducing the available nectar and water for adults. The direct impacts of drought can be mitigated for these species by watering the grass during prolonged or intense droughts.
- Coastal development will also increase habitat loss. Planning controls are possible to minimise this impact.
- A small amount of Floyd's Grass is impacted by cattle grazing and trampling, which also makes it unsuitable for the butterfly.
- Trampling by people, including campers, damages the grass, but can be avoided with access restrictions
- Habitat loss for Floyd's grass (in terms of net habitat) can be mitigate via translocation, but this is last resort.
- Climate. The condition of the grass varies seasonally, annually or at longer time scales depending on temperature, rainfall, drought, flood, etc.

*Additional consideration for the Black Grass-dart Butterfly*

- Climate. Butterfly Numbers vary seasonally, annually or at longer time scales depending on temperature, rainfall, drought, flood, etc.
- inbreeding, disease, predation, and parasitism of larvae and eggs, are known problems for lepidoptera generally, however the level of impact is unknown for this species.
- Demographic stochasticity. This is an issue in very small patches where the timing of adult butterfly emergence and locating mates, etc, relies partly on chance.

#### 2.4 Conceptual model

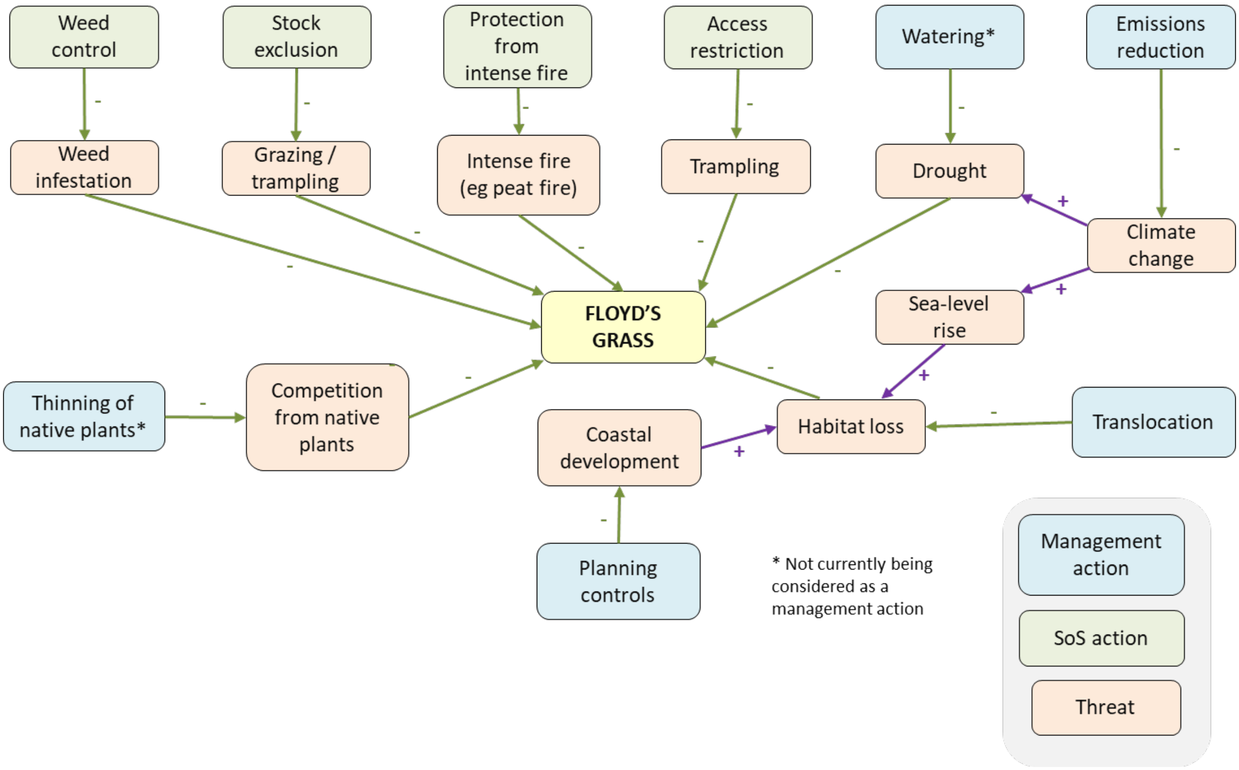


**Fig. 3:** Conceptual model for Floyd's grass

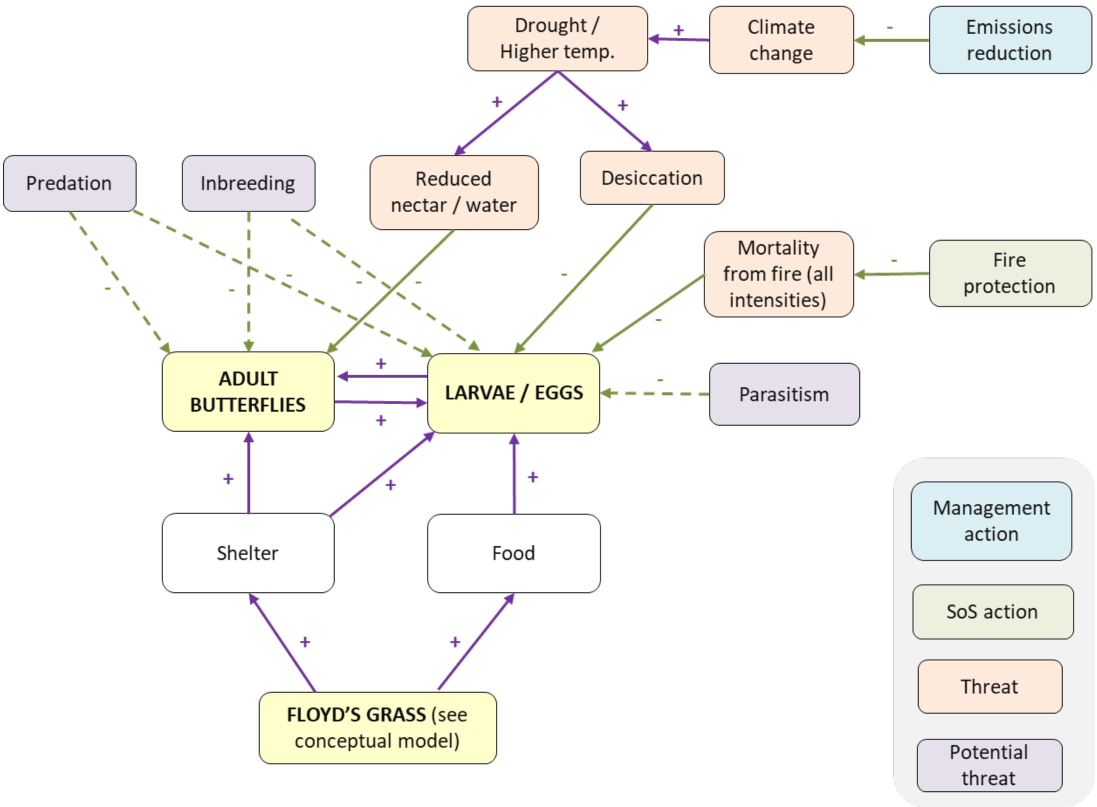


Fig. 4: Conceptual model for Black Grass-dart Butterfly

Mayfield et. al 2025: *Evaluating a structured expert elicitation approach for adaptive conservation: Lessons from five years in practice.*
